# Detection of pre-existing humoral immunity against influenza virus H5N1 clade 2.3.4.4b in unexposed individuals

**DOI:** 10.1101/2025.01.22.634277

**Authors:** Katharina Daniel, Leon Ullrich, Denis Ruchnewitz, Matthijs Meijers, Nico Joel Halwe, Ursula Wild, Jan Eberhardt, Jacob Schön, Ricarda Stumpf, Maike Schlotz, Marie Wunsch, Luana Girao Lessa, El-Sayed Mohammed Abd El-Whab, Maryna Kuryshko, Christopher Dietrich, Andreas Pinger, Anna-Lena Schumacher, Maximilian Germer, Malena Rohde, Christian Kukat, Lutz Gieselmann, Henning Gruell, Donata Hoffmann, Martin Beer, Thomas Erren, Michael Lässig, Christoph Kreer, Florian Klein

## Abstract

The spill-over of Influenza A virus H5N1 clade 2.3.4.4b from cattle to humans highlights the risk of a human H5N1 pandemic. Given the impact of pre-existing immunity on the course and severity of viral infections, we comprehensively assessed the humoral immunity against the H5N1 A/Texas/37/2024 isolate in H5N1-naïve individuals. To this end, we performed complementary binding and neutralization assays on 66 subjects and ranked activities among a panel of 76 influenza A virus isolates. We detected low but distinct cross-neutralizing titers against A/Texas/37/2024 with a 3.9 to 15.6-fold reduction compared to selected H1N1 or H3N2 strains. By cloning and evaluating 136 monoclonal antibodies from memory B cells, we identified potent A/Texas/37/2024-neutralizing monoclonal antibodies in five out of six investigated individuals. These antibodies cross-neutralize H1, compete with antibodies targeting the HA stem, and protect mice from lethal H5N1 challenge. Our findings demonstrate partial pre-existing humoral immunity to A/Texas/37/2024 in H5N1-naïve individuals.

## Main Text

Sporadic human infections with highly pathogenic avian influenza A virus (HPAIV) H5N1 were first documented by spillover from birds to humans in 1997, followed by repeated outbreaks among birds with a total of 954 human cases leading to 464 deaths (48.6% case-fatality rate) between 2003 and 2024.^1^ In March 2024, the first suspected case of H5N1 clade 2.3.4.4b transmission from infected cattle to humans was reported.^2^ This H5N1 variant emerged in wild birds between 2020 and 2022^3^ and was demonstrated to be most likely transmitted to cattle by the end of 2023.^4^ By beginning of January 2025, 66 documented human H5N1 cases had been recorded in the United States, 40 of which were linked to dairy cows.^5,6^ However, as a recent study found serological evidence of past H5N1 infection in 7% of tested farmworkers, the actual case number is likely higher.^7^ The ongoing outbreak of H5N1 infection in cattle increases the risk of more frequent spillover events, further reassortments, and stronger adaptation to human hosts.^8^ Indeed, sequencing analyses have revealed mutations that are related to mammalian host adaptation (including PB2 E627K, M631L, and D701N).^2,4^

As demonstrated by previous influenza pandemics, and more recently by Covid-19, course and severity of disease are strongly influenced by the presence or absence of pre-existing cellular and humoral immune responses. Antibodies against influenza viruses primarily target hemagglutinin (HA) and neuraminidase (NA).^9^ To date, 19 different influenza A virus HA subtypes have been described from which H1, H2, and H3 have been identified to endemically circulate in humans, while avian influenza A virus strains that transmitted to humans mainly belong to subtypes H5, H7, and H9.^10,11^ HA-neutralizing antibody titers as measured by the hemagglutination inhibition (HAI) assay, are currently used as a correlate of protection against H1N1 and H3N2 infections.^12–14^ Moreover, HA-specific monoclonal antibodies (mAbs) have been shown to effectively neutralize various influenza virus strains including different H5N1 isolates.^15–22^

Given the extensive spread of H5N1 clade 2.3.4.4b virus in cattle and the continuous spill-over to humans, there is an urgent need to assess the level of pre-existing humoral immunity in humans. While molecular information on the presence of H5-reactive antibodies in H5N1 unexposed individuals is still limited,^19,23–31^ a detailed understanding of the specific human antibody responses against H5N1 clade 2.3.4.4b is of critical need.

In our study, we determine HA-specific humoral immunity to the recently transmitted human H5N1 isolate A/Texas/37/2024 (clade 2.3.4.4b, genotype B3.13) in H5N1 unexposed individuals. To this end, we apply specific binding (i.e., HA-ELISA and cell-based HA binding FACS) and neutralization (pseudovirus and authentic virus) assays. We place our findings in the broader context of additional 76 human and avian influenza A virus strains and perform deep single B cell and antibody analyses in five H5N1-unexposed and one H5N1-vaccinated individuals. With our study we provide critical knowledge on pre-existing immunity that may inform on potential preventive and therapeutic strategies against the emergent H5N1 influenza virus clade 2.3.4.4b.

### Detectable IgG binding and serum neutralization against H5N1 A/Texas/37/2024 hemagglutinin in H5N1-naïve individuals

To assess potential pre-existing H5N1 clade 2.3.4.4b (A/Texas/37/2024) humoral immunity, we investigated blood samples collected between August 2023 and June 2024 from a regional German cohort of 66 individuals without known exposure to H5N1. All samples were analyzed for HA reactivity and neutralizing activity in three complementary assays (Figure 1A).

**Figure 1.**
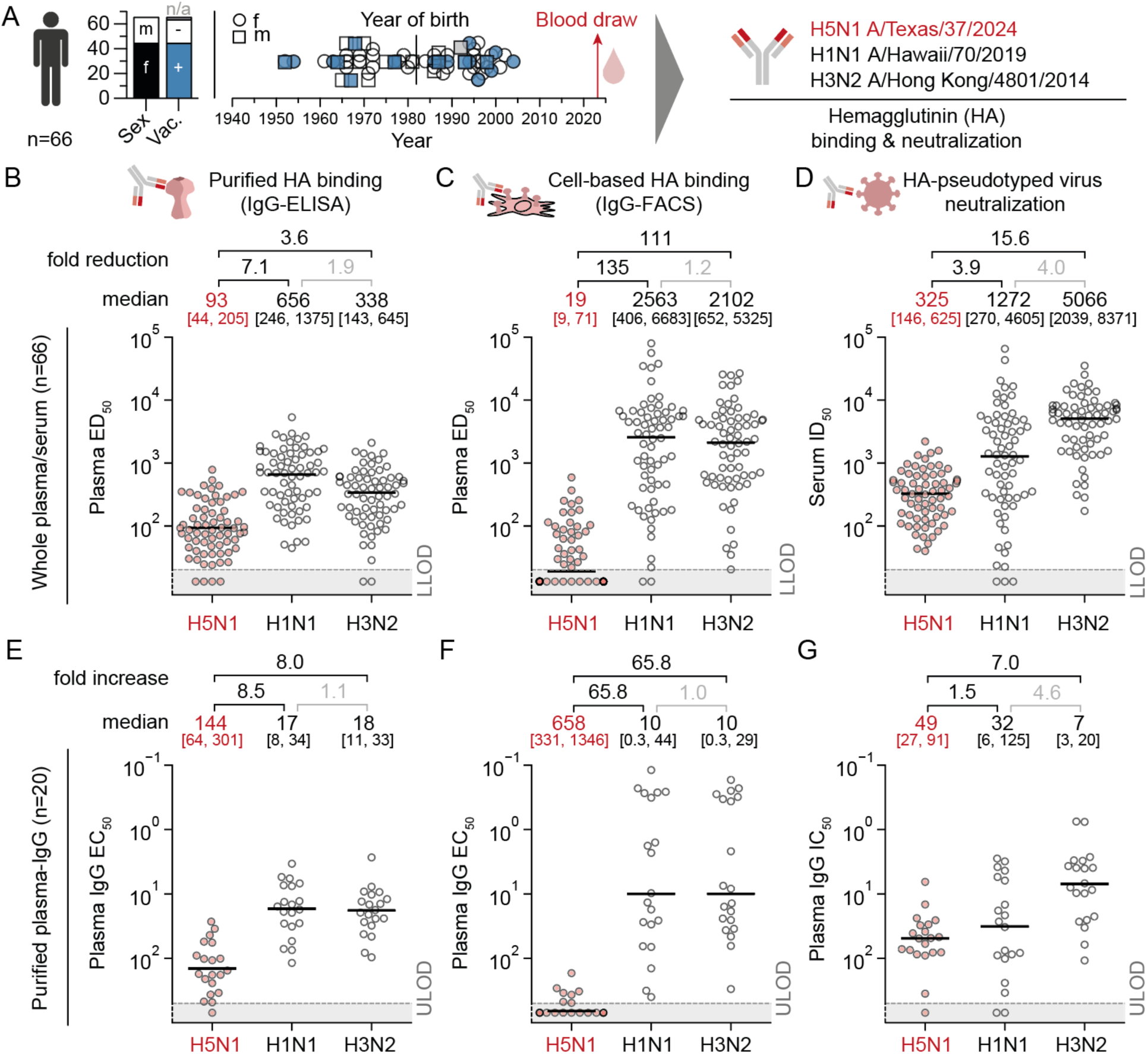
Detection of plasma-IgG binding and serum neutralization against A/Texas/37/2024 HA in H5-naïve individuals. (**A**) Study design and cohort information including sex, vaccination (vac.) status (see Table S1), and year of birth of n = 66 individuals that donated blood between August 2023 and June 2024, as well as the selected strains to test for HA binding and neutralizing activity. (**B**) Half-maximal effective dilution (ED_50_)-values from plasma samples of n = 66 individuals, determined by IgG enzyme-linked immunosorbent assay (ELISA) against recombinant HA proteins derived from the strains in (A). (**C**) ED_50_-values from plasma samples of n = 66 individuals, determined by flow-cytometric measurement of IgG positive cells expressing HA proteins derived from the strains in (A) on their cell surface (see method section for details). (**D**) Half-maximal inhibitory dilution (ID_50_)-values from serum samples of n = 66 individuals, determined by neutralization tests against luciferase-encoding lentivirus pseudotyped with HA variants from the strains in (A). In (B-D) the lowest tested dilution (considered as the lower limit of detection, LLOD) was a 20-fold dilution and is depicted as a dashed line in the plots. The highest tested dilution was 1:327,680. Calculated values below the LLOD were arbitrarily plotted at a constant value below the LLOD for visualization. The pairwise fold reduction in (B-D) is reported with respect to the higher geometric mean. (**E**) Half-maximal effective concentration (EC_50_)-values from purified plasma poly-IgGs for a subset of n = 20 individuals, determined by IgG ELISA as in (B). (**F**) EC_50_-values from purified plasma poly-IgGs for a subset of n = 20 individuals, determined by flow-cytometric measurement of IgG positive cells expressing HA proteins as in (C). (**G**) Half-maximal inhibitory concentration (IC_50_)-values from purified plasma poly-IgGs for a subset of n = 20 individuals, determined by neutralization tests against luciferase-encoding HA-pseudotyped lentivirus as in (D). In (E-G) the highest tested concentration (considered as the upper limit of detection, ULOD) was 500 µg/ml and is depicted as a dashed grey line in the plots. LLOD was 0.0064 µg/ml. Calculated values above the ULOD were arbitrarily set to a constant value above the ULOD for visualization. Y-axes in (E-G) are inverted. The pairwise fold increase in (E-G) is reported with respect to the lower median. Horizontal bars in (B-G) represent the error-corrected median (see methods section for details), which is also reported on top of the graphs together with the 25^th^ and 75^th^ percentiles in brackets. See also Figures S1, S2, S3, S4, S5, S6, and Table S1.

First, we tested human plasma samples in an IgG-ELISA against recombinantly expressed HA proteins (without transmembrane and C-terminal domains) of the A/Texas/37/2024 isolate and two seasonal influenza A viruses as controls (A/Hawaii/70/2019 H1N1 and A/Hong Kong/4801/2014 H3N2; Figure S1; Figure 1B). In 62 out of 66 individuals, we detected H5-reactive IgGs at the lowest plasma dilution of 1:20 with a median half-maximal effective dilution (ED_50_) of 93. This reactivity was substantially lower compared to the 2019 H1N1 and 2014 H3N2 controls (median ED_50_ values of 656 and 338, respectively), reflecting a 7.1- and 3.6-fold decrease, respectively. In addition, we expressed the HA variants as full proteins on the surface of HEK293-6E cells and measured bound plasma-IgGs by flow cytometry (Figure S2, Figure 1C). As a result, median ED_50_ was determined to be 19 for H5N1 (for calculation of values below the limit of detection see Methods). In comparison, median ED_50_ for the controls H1N1 and H3N2 were 2,563 and 2,102, respectively, reflecting a 135 (H1N1) and 111 (H3N2)-fold reduction in binding to cell-surface-expressed HA. Of note, in contrast to determining IgG-binding by ELISA, 37 samples in the cell-based binding assay showed calculated plasma ED_50_ values below the lowest dilution of 1:20.

Next, we assessed potential H5N1 clade 2.3.4.4b neutralizing activity using an HA-pseudotyped lentivirus neutralization assay on Madin-Darby canine kidney cells that overexpress the α2,6-sialyltransferase ST6Gal1^32^ (MDCK-SIAT1; Figure 1D). The median half-maximal inhibitory dilution (ID_50_) for serum was measured to be 325, showing a reduction of neutralizing activity compared to the controls by 3.9-fold (H1N1, ID_50_ of 1,272) and 15.6-fold (H3N2, ID_50_ of 5,066), respectively. However, despite significantly lower neutralizing activity against H5N1 than against H1N1 and H3N2, our findings revealed detectable serum neutralization against H5N1 in all 66 study participants. To rule out unspecific serum inhibition we determined the contribution of IgGs in plasma samples to bind and neutralize H5N1 clade 2.3.4.4b. To this end, we purified poly-IgGs from plasma for a subset of n = 20 study participants (10 male and 10 female), selected to represent a broad age spectrum of the cohort, and tested those in the same three assays for binding (Figure 1E, F) and neutralization (Figure 1G). Fold-reductions measured by half-maximal effective and inhibitory concentrations (EC_50_, IC_50_) of poly-IgGs were similar to plasma/serum samples results: 8.5 (H1N1) and 8.0 (H3N2) for ELISA binding, 65.8 (H1N1) and 65.8 (H3N2) for surface-expressed HA-binding, and 1.5 (H1N1) and 7.0 (H3N2) for HA-pseudovirus neutralization (Figure 1E-G). The predominant role of IgG for H5N1 binding and neutralizing activity was further underlined by a strong correlation between serum and poly-IgG neutralization across all tested strains (Figure S4). Furthermore, we validated the specific effects of IgG antibodies by testing neutralizing activity of plasma samples with and without IgG depletion (Figure S5). IgG depletion led to an almost complete loss of neutralizing activity against H5N1 highlighting the role of IgGs for the serum neutralization against H5. Moreover, testing HA-reactive antibody-depleted poly-IgGs showed a lack of neutralization activity against H5, H1, and H3, but had no impact on SARS-CoV-2 neutralization, demonstrating the specific neutralizing activity of HA-reactive IgGs (Figure S6).

While sex had no impact on serum neutralization (Figure S3A), we detected a significant (p=0.004) 7.1-fold reduction in the neutralization of H1N1 for unvaccinated versus vaccinated individuals (Figure S3B, see Table S1 for self-reported vaccination information). For H5N1, this reduction was less pronounced (2.1-fold; p=0.026). Pseudovirus-based serum neutralization correlated inversely with age for H3N2 (r_p_=0.450, p<0.001) and, to a lesser degree, for H1N1 (r_p_=0.255, p=0.038), but not for H5N1 (r_p_=-0.107, p=0.394; Figure S3C).

We conclude that the majority of individuals without known exposure to H5N1 carry both detectable levels of H5-reactive as well as neutralizing IgG antibodies against H5N1 A/Texas/37/2024.

### H5N1 clade 2.3.4.4b virus-neutralization by H5-naïve individuals’ sera is low in comparison to H5N1-infected cows

To benchmark the potency of the detected H5N1 clade 2.3.4.4b virus neutralization, we tested sera from cows (Figure 2A) that had been experimentally infected with HPAIV H5N1 clade 2.3.4.4b genotypes B3.13 (A/dairy cow/Texas/063224-24-1/2024), which has the same HA sequence as the human isolate A/Texas/37/2024 or DE23-11-N1.3 euDG (A/wild goose/Germany-NW/2024AI02730/2024)^33^, which has seven amino acid differences in comparison to A/Texas/37/2024 HA (one in the signal peptide, four in the head and two in the stalk domain; Figure 2A).

**Figure 2.**
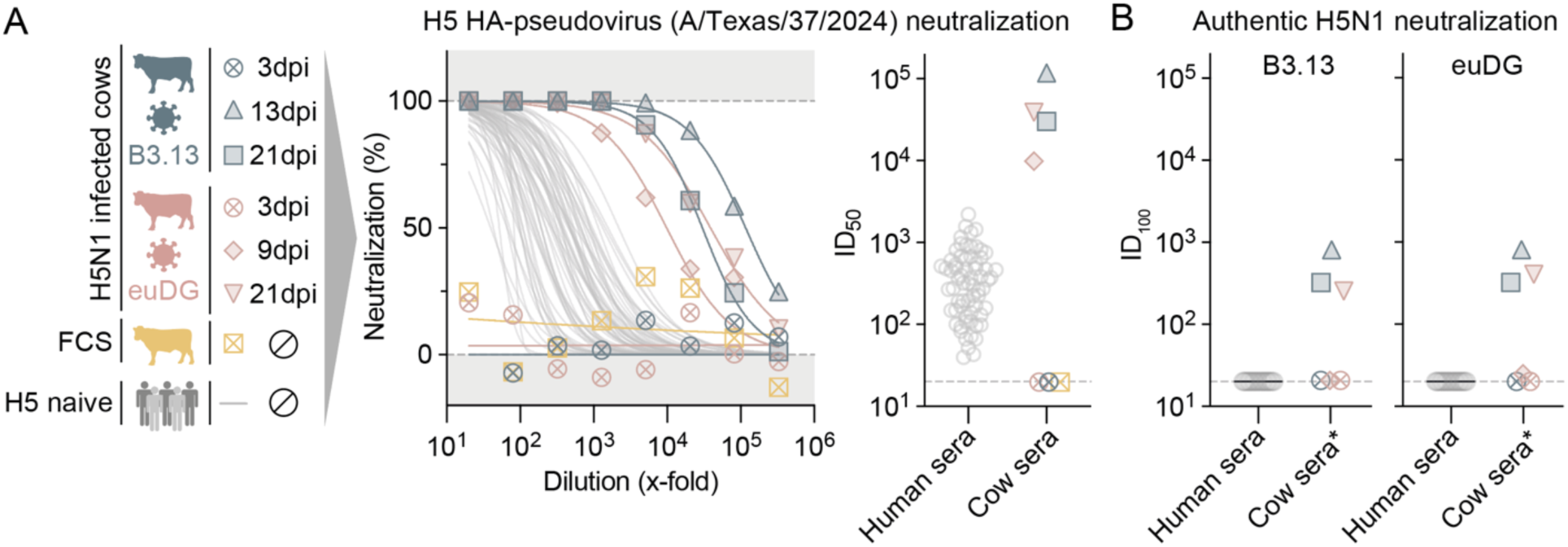
Human H5 pseudovirus and authentic virus serum-neutralization in comparison to H5N1 infected cows. (**A**) Serum samples from cows experimentally infected with H5N1 (H5N1 clade 2.3.4.4b genotypes B3.13 or DE23-11-N1.3 euDG (euDG)), commercial fetal calf serum (FCS), and sera of n = 66 H5-naïve individuals were tested for neutralization activity against pseudotyped lentivirus expressing the HA of H5N1 A/Texas/37/2024. Normalized neutralization from n = 2 technical replicates is depicted by symbols with lines showing non-linear curve fits. Scatter plot shows calculated half-maximal inhibitory dilution (ID_50_, x-fold dilution) for each of the human and bovine serum samples. (**B**) Complete (100%) inhibitory dilutions (ID_100_) were determined for sera of n = 20 H5-naïve individuals and cows experimentally infected with H5N1 by virus neutralization tests against authentic H5N1 B3.13 or euDG. *Data for cow serum neutralization against authentic B3.13 and euDG was taken from Halwe et al., Nature 2024.

While sera from seronegative cows (3 days post H5N1 infection) and commercial fetal calf serum (FCS) did not show any neutralizing activity against H5 A/Texas/37/2024 pseudovirus, we detected potent neutralization with ID_50_ values between 9,843 and 116,984 for cow sera at 9, 13, or 21 days post infection (Figure 2A). In contrast, human sample ID_50_ values were in the range of 40 to 2,199 (Figure 1D and 2A). In addition, 20 human and 6 cow samples^33^ were tested against authentic HPAIV H5N1 clade 2.3.4.4b genotypes B3.13 (A/dairy cow/Texas/063224-24-1/2024) and euDG (A/wild goose/Germany-NW/2024AI02730/2024) in a microneutralization test on MDCK II cells. While authentic virus neutralization was detectable for cow sera with complete (100%) inhibitory dilution (ID_100_) ranging from 25 to 813,^33^ we were not able to detect authentic H5N1 B3.13 or euDG neutralization in the 20 human sera (Figure 2B). When comparing pseudovirus ID_50_ and authentic virus ID_100_ from four infected cows with H5N1 antibody responses, the H5 ID_50_ values determined by pseudovirus were about 25-fold higher than ID_100_ values from authentic virus neutralization tests. Similarly, we found strong correlation between ID_50_ and ID_100_ values from control H1N1 pseudovirus and wild-type neutralization assays (Figure S8).

We conclude that H5N1 clade 2.3.4.4b serum neutralization in H5N1-naïve individuals is substantially lower than in cows that mounted an H5N1-specific antibody response. Moreover, we conclude that the performed authentic virus neutralization assay is less sensitive and does not detect very low levels of neutralizing antibodies in human sera.

### Ranking serum-neutralizing activities against H5N1 A/Texas/37/2024 and recent and historical Influenza A virus isolates

In order to compare H5N1 A/Texas/37/2024 neutralization with a broader number of recent and historical influenza A virus strains, we compiled and analyzed 83,118 influenza virus sequences from HA subtypes H1-H18 and built a global phylogenetic tree (Figure 3A). Next, we selected and produced 77 different HA-pseudotyped virus variants of subtypes H1, H2, H3, H5, H7, and H9 (Table S2) that (i) were geographically spread, (ii) included frequent and pre-dominant mutations of particular lineages, and (iii) have been associated with human infections (Figure 3A; see methods section for details).

**Figure 3.**
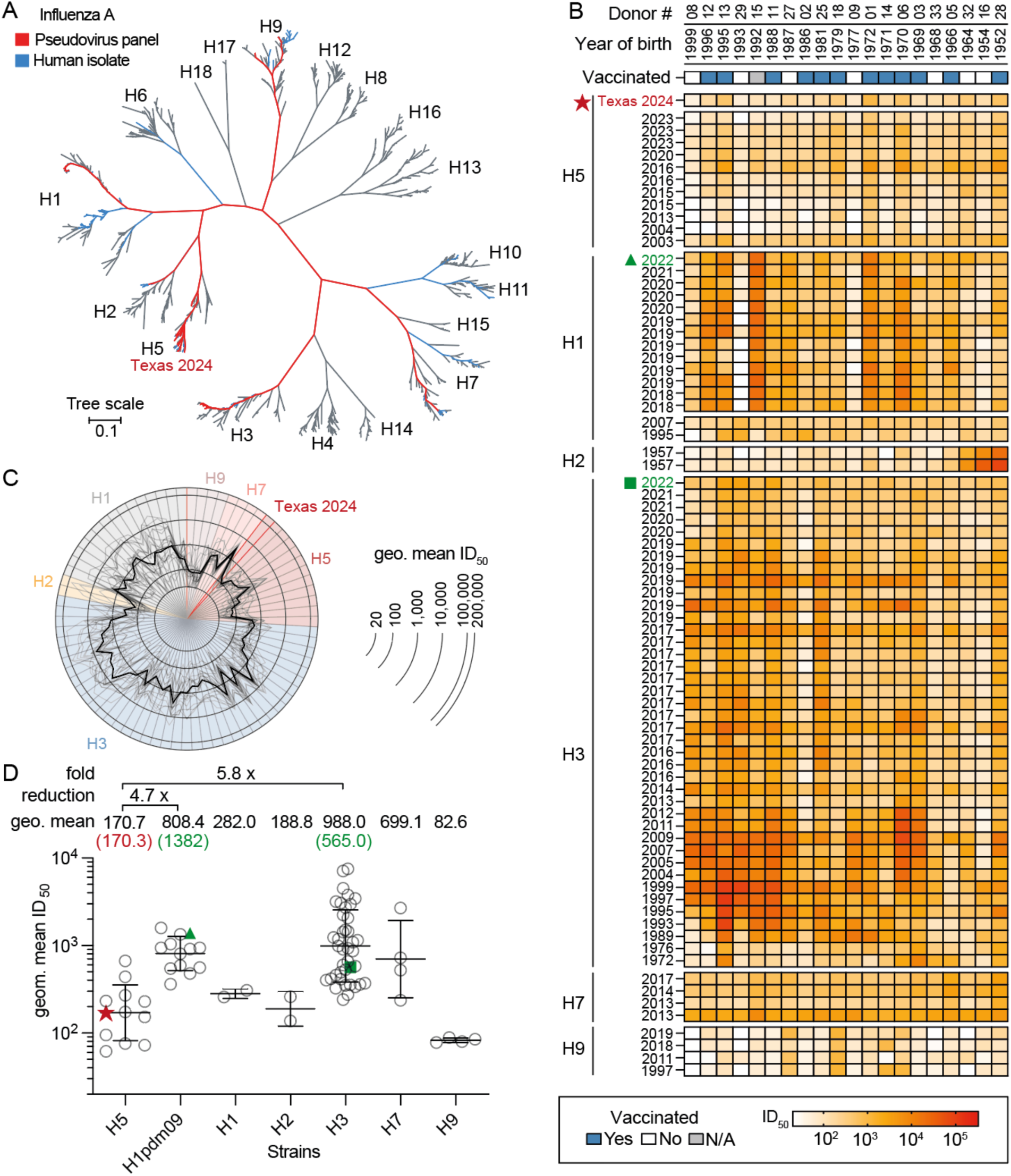
Comparison of serum neutralization across seasonal and avian influenza HA-pseudotyped lentiviruses. (**A**) Phylogenetic tree of HA variants based on 83,118 quality-controlled sequences. 77 strains covering H1, H2, H3, H5, H7 and H9, which were all isolated from humans were selected based on geography and frequency to include critical mutations of the lineages for pseudovirus generation. (**B**) Half-maximal inhibitory dilution (ID_50_) of 20 sera tested by HA-pseudotyped lentivirus neutralization against the 77 selected strains from (A). A/Texas/37/2024 is highlighted by a red star. The most recent H1 and H3 strains are highlighted by a green triangle and square, respectively (**C**) Comparison of geometric mean ID_50_ values for each strain. Each virus strain is represented by one pie slice. Light gray lines show ID_50_ values from each individual (i.e., columns in (B)). The black line shows the geometric mean ID_50_ over all individuals. (**D**) HA-groupwise comparison of geometric mean ID_50_s. Each symbol represents the geometric mean ID_50_ over all 20 individuals for a given strain (i.e., the row geometric mean for each strain in (B)). A/Texas/37/2024 is indicated by a red star. The most recent H1 and H3 strains are indicated by a green triangle and square, respectively. Geometric mean ID_50_ values for these strains are reported in the respective color in brackets above the graphs. Average lines depict geometric means over all strains from a subtype, which are reported above the graphs. Error bars depict geometric standard deviation. The fold reduction of the geometric mean for H5 vs. H1pdm09 and H3 is depicted in the graph. See also Figure S7 and Table S2.

When testing serum from 20 of the 66 study participants against the 77 pseudoviruses panel (Figure 3B), we detected neutralizing activities ranging from ID_50_ values of ≤ 20 up to 111,102 (Figure 3B and C). When calculating the geometric mean per individual (i.e. column-wise), we found higher H1 neutralizing activity in individuals that recently underwent influenza vaccination (1174, vs. 194.3; p=0.001 for log_10_-transformed geometric mean ID_50_ values; Figure 3B, Table S1, Figure S7A and B). Moreover, H2 and H3 neutralization was associated with year of birth: Higher H2 neutralizing activity was detected in individuals that were born when H2N2 was circulating (1957-1968) and higher H3 neutralization in younger individuals that were born after the emergence of H3N2 (1968; Figure 3B, Figure S7A).

To rank the neutralizing activity against H5N1 A/Texas/37/2024, we calculated the geometric mean over all 20 individuals for each strain (Figure 3C) and grouped them by their HA subtype (Figure 3D). Within the tested H5 strains, the geometric mean ID_50_ values ranged from 61 (A/Vietnam/1203/2004, clade 1) to 665 (A/Hubei/29578/2016, clade 2.3.4.4h), with the H5N1 A/Texas/37/2024 (ID_50_=170.3) being close to the group’s geometric mean of 170.7 (Figure 3D) and significantly higher than A/Vietnam/1203/2004 (p<0.001, two-tailed paired t-test; Figure S7C). In comparison, neutralizing serum activities against H1pdm09 (808.4) and H3 (988.0) were significantly higher (adjusted p=0.003 and p<0.001, respectively, tested by Kruskal-Wallis with Dunn’s correction for multiple testing; Figure 3D).

In conclusion, comprehensive mapping revealed that neutralizing serum activity against H5N1 A/Texas/37/2024 was near average among all tested H5 strains, but significantly lower compared to influenza strains of the subtypes H1 (2.1 to 9.4-fold) and H3 (1.4 to 44-fold).

### Detection of memory B cells that encode for H5N1 A/Texas/37/2024 HA-neutralizing antibodies in H5-naïve individuals

Next, we investigated the H5N1 A/Texas/37/2024 humoral immune response on a molecular level. To this end, we adapted previous strategies to isolate HA-specific memory B cells^34,35^ by introducing the Y98F mutation into the A/Texas/37/2024 hemagglutinin that reduces HA-binding to sialic acid.^36^ Using this protein as a bait, we single-cell sorted IgG^+^ memory B cells from six H5-naïve individuals with A/Texas/37/2024 serum-ID_50_ values between 43 and 2,199 as well as one individual that had received an experimental H5N1 vaccine in 2008/2009 (clade 1, A/Vietnam/1203/2004; Figure 4A, Figure S9; ID_50_=2,343).

**Figure 4.**
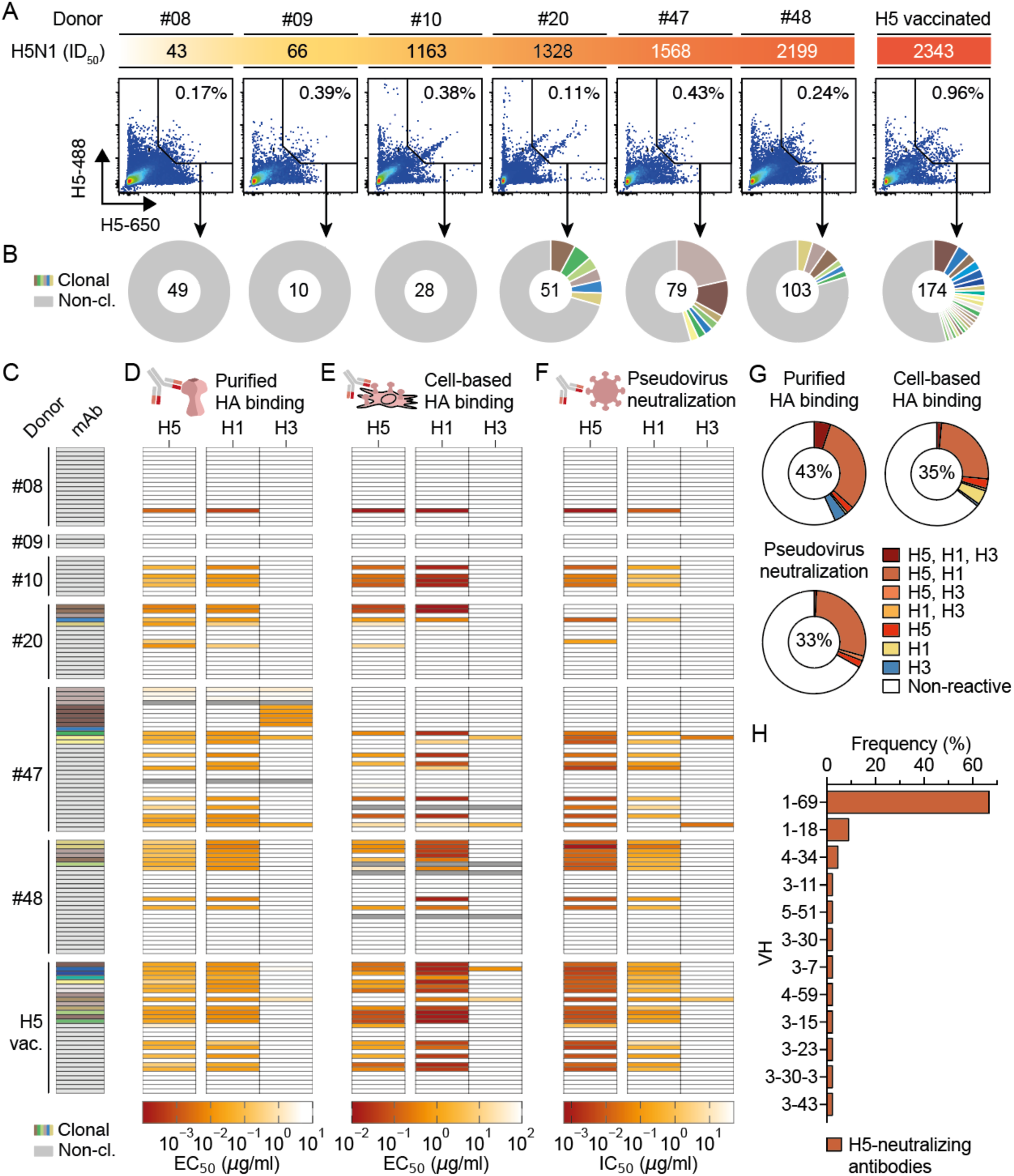
Isolation of HA A/Texas/37/2024-reactive memory B cells. (**A**) Dot plots of IgG^+^ B cells positive for H5N1 TEXAS/37/2024 HA-Y98F labeled with different DyLight488 or DyLight650 from n = 6 H5-naïve (Donor 08, 09, 10, 20, 47 and 48) and one H5-vaccinated individual. Numbers in plots indicate the percentage of double positive H5-reactive B-cells within IgG^+^ B cells. The half-maximal inhibitory dilution (ID_50_, x-fold dilution) against pseudotyped lentivirus expressing the HA of H5N1 A/Texas/37/2024 is depicted above the plots. (**B**) Clonal relationship of H5-reactive B cells. Total number of analyzed sequences is indicated in the center of the pie charts. (**C-F**) 136 antibodies isolated from individuals presented in (A) were analyzed for their binding and neutralization against A/Texas/37/2024 (H5), A/Hawaii/70/2019 (H1) and A/Hong Kong/4801/2014 (H3) hemagglutinin by IgG enzyme-linked immunosorbent assay (D), cell-based binding assay (E) and pseudovirus neutralization assay (F). Heatmaps show half-maximal effective/inhibitory dilution (ED_50_/ID_50_)-values. Clonal relationship of tested antibodies is illustrated in (C). (**G**) Pie charts summarizing the reactivity of tested antibodies for the three assay systems used (D-F). Numbers in plots indicate the percentage of HA-reactive antibodies. (**H**) Frequencies of heavy chain V-gene usage from n = 54 H5-binding and/or neutralizing antibodies. See also Figure S9.

By performing B cell receptor (BCR) RT-PCR,^37^ we amplified a total of 494 functional immunoglobulin heavy chains. Sequence analysis revealed the presence of clonally related memory B cells in 4 out of 7 individuals (Figure 4B). From all amplified and sequenced BCRs, we cloned and produced 136 monoclonal antibodies (Figure 4C) and tested them for binding against recombinant HA protein and cell-associated HA, as well as determined virus neutralizing activity against H5N1 (A/Texas/37/2024), H1N1 (A/Hawaii/70/2019), and H3N2 (A/Hong Kong/4801/2014) hemagglutinin pseudoviruses (Figure 4D-F). Notably, we identified H5N1 A/Texas/37/2024 binding and neutralizing antibodies in 5 out of 6 H5N1-naïve individuals as well as in the H5N1 vaccinated subject (Figure 4D-F). In the H5N1-naïve subject #9 with low A/Texas/37/2024 virus neutralization, no H5 reactive monoclonal antibodies (mAbs) were isolated. However, for this individual only 3 antibodies were tested.

For each of the 5 H5N1-naïve individuals in whom we detected A/Texas/37/2024-reactive antibodies, 1 to 14 mAbs bound to recombinant H5 protein, 1 to 7 bound to cell-associated H5, and 1 to 10 neutralized H5-pseudotyped virus. For the H5N1 vaccinated individual, 18 out of 30 antibodies showed H5 binding and neutralization. Overall, there is a strong overlap for binding and neutralizing activity across the three assays with in total 43%, 35%, and 33% of monoclonal antibodies tested showing reactivity with recombinant HA, cell-surface-expressed HA, and HA-expressing lentivirus, respectively (Figure 4G). Moreover, our data demonstrated that the great majority of H5-reactive antibodies are cross-binding (90 to 92%) as well as cross-neutralizing H1 (89%; Figure 4D-G). In contrast, cross-binding and -neutralization of H3 was only detected in 5 to 13% and 7%, respectively (Figure 4D-G). Finally, sequence analysis revealed that > 60% of all H5-neutralizing antibodies utilize V_H_ gene segment 1-69 (Figure 4H).

We conclude that H5N1-naïve individuals carry H5-reactive memory B cells encoding for A/Texas/37/2024 neutralizing antibodies. We further conclude that the majority of these antibodies are cross-reactive with H1 and preferably facilitate V_H_1-69.

### Antibodies from H5-naïve individuals can be highly neutralizing, predominantly compete with HA stem antibodies, and are protective in vivo

To characterize the potency of the isolated antibodies, we compared 45 isolated H5-neutralizing antibodies with previously published influenza virus antibodies. To this end, we produced a reference panel of nine HA-reactive antibodies (seven of which have been documented as H5-reactive before, Table S3),^15–18,38–42^ including five antibodies that have been in clinical development (MEDI8852, 39.29, CR9114, CR6261 and CR8020).^15–17,42^ All reference antibodies were tested for surface-expressed HA-binding (Figure S10A) as well as pseudovirus neutralization against three H1, one H2, three H3, and five H5 strains (including A/Texas/37/2024; Figure S10B). We found that six out of the seven previously described H5-reactive antibodies retained binding and neutralizing activity against the H5N1 A/Texas/37/2024 isolate with neutralizing potencies in IC_50_ ranging from 0.003 to 0.97 µg/ml (Figure S10B). Of note, the monoclonal antibody FluA-20, which was previously described to bind but not neutralize group 1 and 2 HA subtypes *in vitro,*^43,44^ bound to A/Texas/37/2024 HA, but did not neutralize the corresponding pseudovirus, highlighting the specificity of the pseudovirus neutralization assay (Figure S10C).

Next, we compared the H5N1 A/Texas/37/2024 neutralizing activity of our 45 isolated H5-neutralizing antibodies with the neutralizing reference antibodies. Notably, 67% to 100% of tested antibodies from the H5N1-naïve subjects #10, #20, #47, and #48, demonstrated neutralizing activity within the range of the reference antibodies (Figure 5A). Moreover, two isolated monoclonal antibodies of subject #8 and #48 demonstrated even higher (IC_50_ of 0.0006 and 0.0008 µg/ml) H5N1 A/Texas/37/2024 activity than the most potent reference antibody (MEDI8852, IC_50_ of 0.003 µg/ml) in the pseudotyped neutralization assay (Figure 5A).

**Figure 5.**
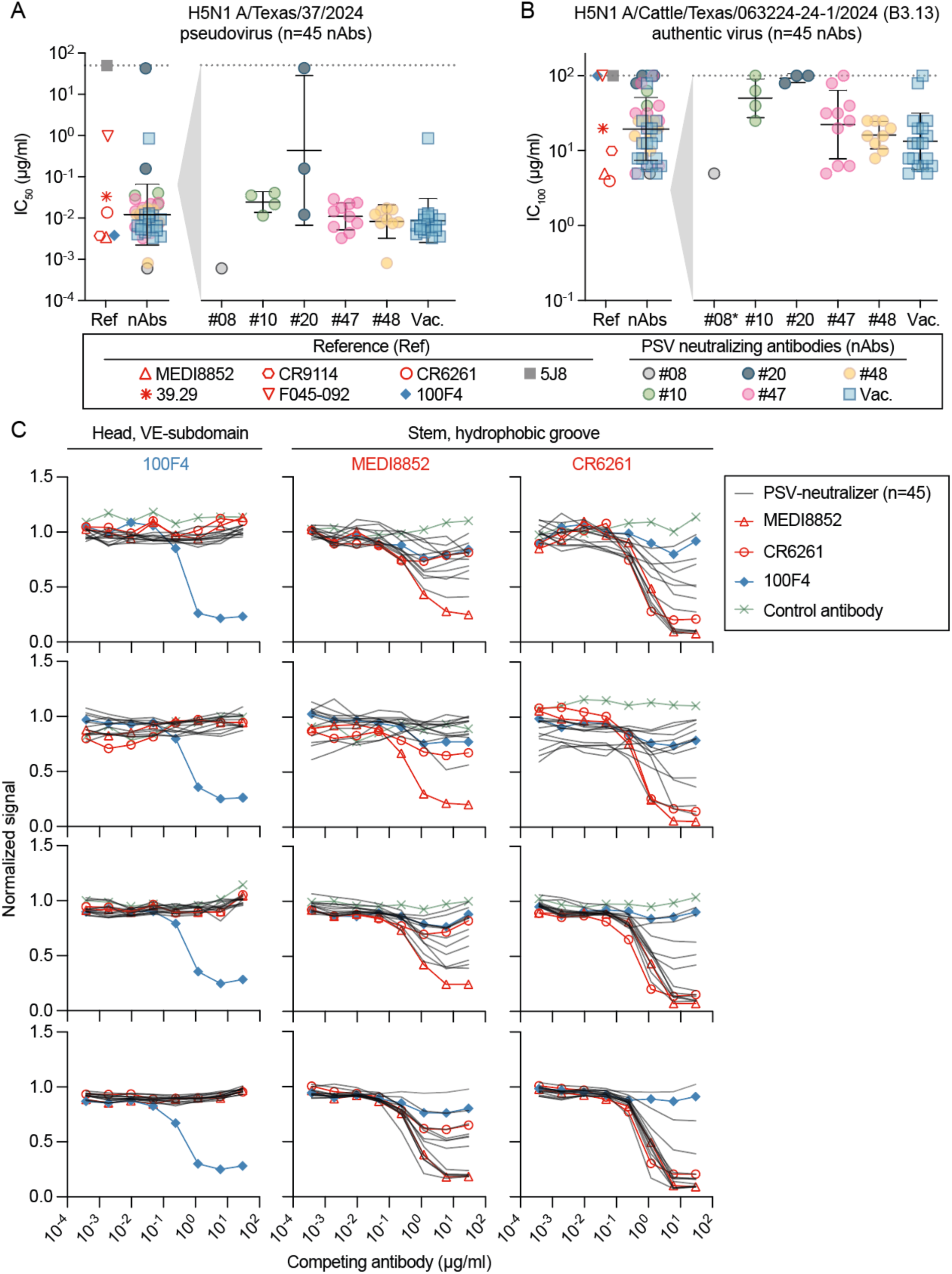
H5-neutralizing antibodies from H5-naïve individuals in comparison to published and clinically developed antibodies. (**A**) Half-maximal inhibitory concentration (IC_50_)-values from n = 45 H5-neutralizing antibodies (nAbs) isolated from H5-naïve or H5-vaccinated individuals (see also figure 4) and n = 7 reference antibodies, determined by neutralization assay against pseudotyped lentivirus expressing the HA of A/Texas/37/2024 (H5N1 B3.13). (**B**) Complete (100%) inhibitory concentrations (IC_100_) from n = 45 H5-neutralizing antibodies (nAbs) isolated from H5-naïve or H5-vaccinated individuals and n = 7 reference antibodies (Ref), determined by microneutralization test against authentic highly pathogenic avian influenza virus (bovine isolate A/dairy cow/Texas/063224-24-1/2024, genotype B3.13). (**C**) Binding of n = 11 H5-pseudovirus (PSV) neutralizing antibodies and n = 3 reference antibodies to recombinant HA protein of A/Texas/37/2024 following incubation with increasing concentrations of indicated biotinylated competitive monoclonal antibodies (100F4, MEDI8852 and CR6261), measured by competitive ELISA. The SARS-CoV-2 antibody CnC2t1p1_D6 was used as a control. Mean values and error bars in (A) and (B) depict geometric mean and geometric standard deviation. See also Figures S10, S11, and Table S3.

To cross-validate neutralizing activity in the authentic virus assay, we tested the 45 H5 pseudovirus-neutralizing and the seven H5-reactive reference antibodies against the bovine isolate A/dairy cow/Texas/063224-24-1/2024 (B3.13) and the related European isolate A/wild goose/Germany-NW/2024AI02730/2024 (euDG) in a microneutralization assay on MDCK II cells. 40 and 36 out of 45 H5 pseudovirus-neutralizing antibodies also neutralized authentic B3.13 and euDG, respectively, while four out of seven H5-reactive reference antibodies neutralized both B3.13, and euDG (Figure 5B, Figure S11A). Isolated and reference antibodies also showed comparable neutralization against H1N1 pseudovirus and authentic virus of isolate A/Sydney/5/2021 (Figure S11B).

To identify possible binding regions of the isolated monoclonal antibodies from naïve individuals, we performed a competitive ELISA between eleven isolated monoclonals (from donors #10, #20, #47, #48, and the H5-vaccinated individual; A/Texas/37/2024-IC_50_ between 0.0008 to 0.02 µg/ml) and two stem-specific reference antibodies (MEDI8852 and CRC6261), as well as one vestigial esterase subdomain (VE)-binding antibody (100F4) on A/Texas/37/2024 recombinant HA protein. We found 39 out of 45 antibodies to compete with biotinylated stem antibodies MEDI8852 and/or CRC6261, while no binding inhibition of the H5-specific VE-subdomain antibody 100F4 to A/Texas/37/2024 HA (Figure 5C) was detected.

Finally, we assessed the capacity of H5-reactive antibodies to protect mice from a lethal H5N1 infection. To this end, we selected two of our most potent antibodies from individuals 48 and 47 (48H5-1-1G9K or 47H5-1-2B12K) and immunized BALB/c mice intraperitoneally (i.p.) with 10 mg/kg of either one of the antibodies or an unrelated SARS-CoV-2 control antibody (DZIF-10c^45,46^), before challenging them with a lethal dose of H5N1 (bovine isolate A/dairy cow/Texas/063224-24-1/2024, genotype B3.13; Figure 6A). Animals were weighed daily and clinically scored for 10 days post infection. All animals pre-treated with the SARS-CoV-2 antibody DZIF-10c died within the 10 days or had to be euthanized due to clinical scoring and severe weight loss (Figure 6B and C). In contrast, all animals receiving the H5-specific antibodies survived the H5N1 infection (Figure 6B), with only one animal in the mAb2 group showing weight loss >10% (Figure 6 C).

**Figure 6.**
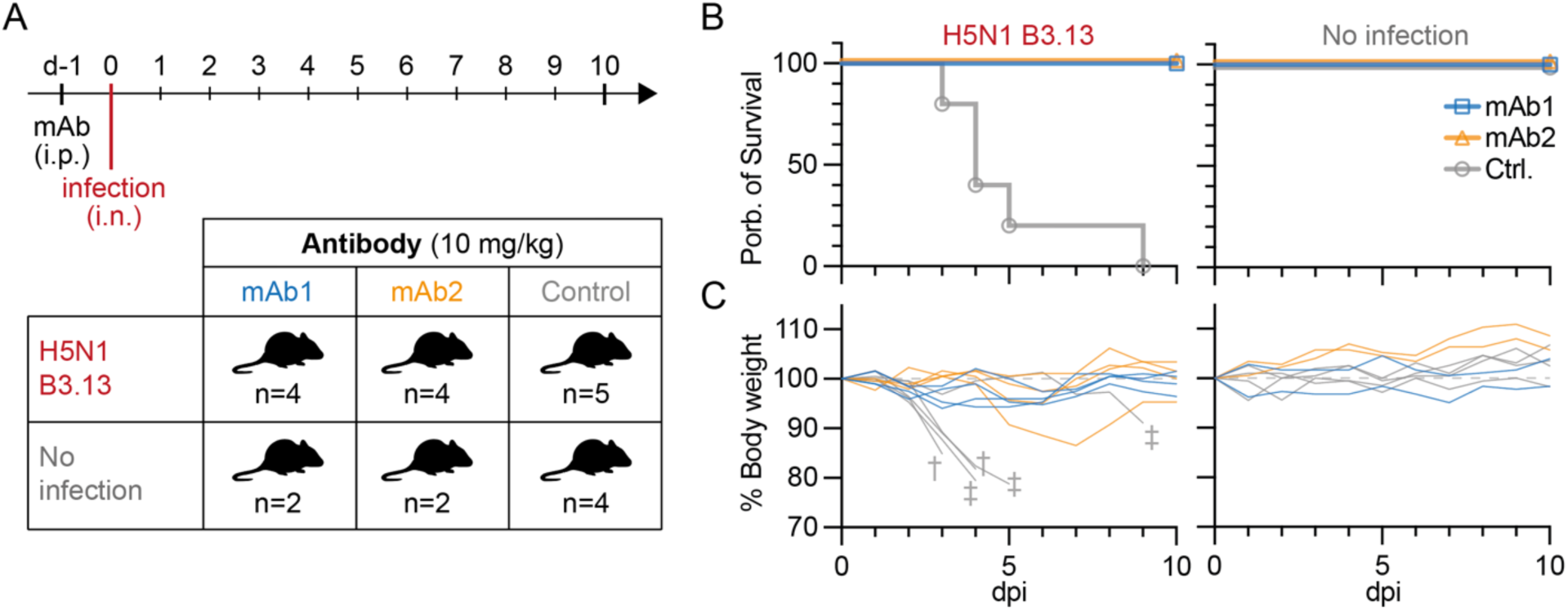
*In vivo* protection of H5-reactive antibodies from lethal H5N1-challenge. (**A**) Experimental outline and antibody selection. Animals were pre-treated intraperitoneal with 10 mg/kg of the H5-reactive antibodies 48H5-1-1G9K (mAb1) or 47H5-1-2B12K (mAb2) and challenged intranasally with a lethal dose of H5N1 (A/dairy cow/Texas/063224-24-1/2024, genotype B3.13) or a mock control (no infection). (**B**) Probability of survival after lethal H5N1 challenge or no infection. (**C**) Body weight loss in relation to body weight at d0 after lethal H5N1 challenge or no infection. † Animals died. ‡ Animals were euthanized due to reaching their individual weight limit or the humane endpoint.

We conclude that H5N1 A/Texas/37/2024 neutralizing monoclonal antibodies from H5-naïve individuals can reach and exceed potencies of clinically developed antibodies, are likely to target the HA-stem region, and can protect mice from lethal influenza H5N1 challenge.

## Discussion

HPAIV H5N1 has sporadically infected humans since 1997.^47^ However, the emerging and uncontrolled outbreak of H5N1 in dairy cattle highlights a novel source of cross-species transmission and increases the risk of viral reassortment and eventually human adaptation. Previous studies have documented cross-binding or cross-neutralizing activity in human sera against earlier H5 isolates^25,48–50^ and rare H5 cross-reactive antibodies have been isolated from H5-unexposed individuals.^15–22,50^ However, human serology studies have produced conflicting results^19,23–26^ and H5 continues to evolve into an antigenically highly divers subtype.^51,52^ This ongoing diversification raises critical questions about the prevalence and potency of pre-existing antibodies, particularly against the most recently emerged H5N1 clade 2.3.4.4b descendants.

Here, we present what is, to our knowledge, the first comprehensive analysis of the humoral immune response to the recent H5N1 clade 2.3.4.4b isolate A/Texas/37/2024, combining complementary binding and neutralization assays at both the polyclonal and monoclonal single-B-cell level. Our results show that the majority of investigated H5N1-naïve individuals (n = 66) harbor detectable H5-reactive antibodies, evidenced by 94% positivity in ELISA, 45% in the cell-based HA-binding assay, and 100% in a highly sensitive pseudovirus neutralization assay. Differences between the three assays likely reflect both a limited correlation between antibody binding strength and neutralization potency, which has also been reported for SARS-CoV-2 antibodies,^53^ as well as the high sensitivity of the pseudovirus assay,^49,54^ enabling detection of neutralizing activity at antibody concentrations below the binding assay detection limits. However, H5-neutralizing ID_50_ values from these 66 individuals were 3.9 and 15.6-fold lower compared to selected H1 or H3 variants, and up to 9.4- and 44-fold lower when measuring a subset of participants against an expanded HA-pseudovirus panel of previously circulating H1N1 and H3N2 isolates. Moreover, the highest detected ID_50_ of H5-naïve human serum was 4.5 to 53.2-fold lower than the serum ID_50_ values from experimentally H5N1 infected cows. Notably, human serum did not exhibit neutralization in an authentic virus microneutralization assay, likely caused by differences in determining the neutralizing activities (ID_100_ vs. ID_50_) as well as the higher sensitivity of the pseudovirus assay.^49,54^ The latter may be attributed to different target cell-lines (MDCK-SIAT1 vs. MDCK II), luminescent vs. microscopic read-out, or different HA densities and accessibilities as discussed elsewhere.^54^ Interestingly, geometric mean neutralization of H5-pseudotyped viruses from different H5 clades and isolates ranged over one log_10_ scale, indicating the impact of antigenic evolution within animal reservoirs on pre-existing immunity. Variations in assay sensitivity and readouts as well as the use of different H5 clades and strains may thus explain publications that report no H5 serum neutralization in the majority of H5N1-naïve individuals.^19,23,24,26–31^

Importantly, we were able to isolate memory B cells encoding for A/Texas/37/2024 HA-binding and neutralizing antibodies from 5 out of 6 H5-naïve individuals and demonstrate that these antibodies can protect mice from lethal H5N1 challenges. The majority of tested antibodies was characterized by H1 cross-reactivity, V_H_ gene segment 1-69 usage, and competition with stem-directed antibodies. The predominance of V_H_1-69 among isolated antibodies is generally consistent with reports on group 1 influenza A virus-reactive antibodies and particularly pronounced within H5 stem-reactive antibodies.^35,55^ In addition, HA-stem-specific antibody levels have been demonstrated to increase upon heterologous H5N1 vaccination,^56,57^ suggesting a critical role of stem-antibodies to mediate H5 cross neuralization. However, further research is required to delineate the complex interplay of immune histories in shaping H5 reactivity and to identify the precise epitopes. Interestingly, recent studies demonstrated the presence of H5N1 cross-reactive neuraminidase-targeting antibodies as well as H5N1 cross-reactive T-cells in H5N1-naïve individuals,^26,58^ suggesting a multifaceted immunity against H5N1 clade 2.3.4.4b virus. In addition, H5-specific vaccines have shown promising results, achieving up to 95% seroconversion against the clade 2.3.4.4b isolate A/Astrakhan/3212/2020.^59^

Monoclonal antibodies represent an additional preventive and therapeutic option, as has been demonstrated for various infectious diseases, such as RSV,^60^ Ebola virus,^61^ or SARS-CoV-2.^62^ As several broadly neutralizing monoclonals against group 1 and/or group 2 influenza viruses have been reported,^15–18,20,39,42,63^ we investigated their H5N1 binding and neutralizing capability and were able to show that the majority of tested broadly-reactive influenza virus directed mAbs (MEDI8852, 39.29, CR9114, CR6261, F045-092, and 100F4) retained potent neutralizing capacity against H5N1 clade 2.3.4.4b *in vitro*.

In summary, we demonstrate detectable but low levels of A/Texas/37/2024 humoral immunity in H5N1-naïve individuals with the presence of memory B cells, which encode for potent H5-neutralizing antibodies that protect BALB/c mice from a lethal H5N1 infection. This is in contrast to the lack of cross-neutralizing antibodies for SARS-CoV-2 before the emergence of the Covid-19 pandemic.^64,65^ However, in our study, we neither infer risks for H5N1 transmission from cattle to humans nor within the human population. Moreover, as we have learned from influenza and other viral pathogens, single mutations can be sufficient to mediate viral escape from humoral immunity as well as from antibody therapeutics.^66–70^ Therefore, further studies are needed to determine reliable correlates of protection for newly emerging influenza virus variants.

### Limitations

This study is subject to the following limitations. First, the cohort is restricted to 66 individuals from the local area of Cologne, Germany and exhibits an above-average vaccination rate of 70%. Second, only a limited number of 6 individuals from this cohort were subjected to single B cell analysis. Third, direct comparison of neutralizing activity between human and avian HA subtypes might be affected by different preferences for α2-6-over α2-3-linked glycans: Human and avian influenza A virus subtypes differ in their specificity for α2-3-linked or α2-6-linked sialic acids, with H5N1 (including A/Texas/37/2024) favoring α2-3-linked sialic acid, and human influenza A viruses favoring α2-6-over α2-3-linked sialic acid.^71–74^ To determine pseudovirus infectivity, we used MDCK-SIAT1 cells, which overexpress α2-6 linked sialic acid^32^ and are thus more readily infected by human subtype HA-pseudotyped viruses (i.e., H1, H2, H3). Fourth, to increase chances of isolating rare H5-reactive antibodies, we used a less stringent sorting gate. This comes at the cost of sorting specificity, indicated by a fraction of about 1/3 of reconstituted antibodies binding to recombinant H5 HA. Finally, our analyses do not include clinical parameters and therefore we cannot draw conclusions on how pre-existing levels of H5-reactive antibodies reduce disease severity and clinical outcomes.

## Materials and Methods

### Cell lines

MDCK-SIAT1 (Merck, cat. no. 5071502) cells were cultured in Minimum Essential Medium Eagle (MEM, Sigma-Aldrich) containing 10% FBS (Sigma Aldrich), 1% Penicillin-Streptomycin (PS, Gibco), 1 mM L-Glutamine (Gibco) and 1% MEM-NEAA (Gibco). HEK293T (ATCC, cat. no. CRL-11268) and HEK293T-ACE2 cells (BEI Resources Catalog# NR-52511) were maintained in DMEM (Gibco) with 10% FBS, 1% PS, 1 mM L-Glutamine and 1 mM Sodium pyruvate (Gibco). All cell lines were grown on 15 cm tissue culture dishes (TPP) at 37°C and 5% CO_2_. For passaging, cells were washed with PBS, detached by incubation with 0.05% Trypsin-EDTA (Gibco) for HEK293T and HEK293T-ACE2 cells and 0.25% Trypsin-EDTA (Sigma-Aldrich) for MDCK-SIAT1 cells and the required number of cells was transferred to a tissue culture dish with fresh growth medium. HEK293-6E cells (National Research Council of Canada, NRC file 11565) were cultured under constant shaking at 110 rpm at 37°C and 6% CO_2_ in FreeStyle 293 Expression Medium (Thermo Fisher) supplemented with 0.2% PS.

### Enrollment of human subjects and study design

All blood samples were collected from study participants who gave their written informed consent under the “EIKIM”-study protocol (DRKS00026266) approved by the Ethics Committee of the University of Cologne (#21-1468).

### Processing of serum, plasma and whole blood samples

Serum was collected by centrifugation using Serum-gel tubes (Sarstedt) and stored at - 80°C. For isolation of plasma and peripheral blood mononuclear cells (PBMCs), blood samples were collected in EDTA tubes (Sarstedt) and centrifuged. Plasma was collected and stored at −80°C. PBMCs were isolated by density gradient centrifugation using Leucosep tubes (Greiner Bio-one) prefilled with Histopaque-1077 (Sigma) according to the manufacturer’s instructions.

### Strain selection

To select Influenza A strains, we generated individual phylogenetic sub-trees for H1, H2, H3, H5, H7, and H9. The trees were partitioned into clades which a) are genetically different, b) reach a significant frequency (20%) in the global population and are geographically wide-spread, and, specifically for H3 and H1(pdm09), c) are antigenically different based on a antigenic inference method^75^ using ferret neutralization data.^76^ From each significant clade, we then selected a representative isolate. For seasonal human influenza, if available, vaccine viruses or isolates used as reference virus in ferret neutralization assays were prioritized. For the subtypes H1(seasonal), H5, H7 and H9, isolates were required to be at least 5 amino acids (aa) mutations different from each other or were recently collected from exposed individuals. If available, human isolates or candidate vaccine viruses (CVVs) were prioritized. For H9, we also selected one animal isolate from Y280A to cover H9 evolution.

### HA cloning

Full HA coding regions (see Table S2 for strain names and EPI numbers) with original leader peptides were codon optimized (Vectorbuilder Codon Optimization Tool) and ordered as gene fragments from Twist Bioscience. Lyophilized gene fragments were solved in nuclease-free water (Ambion) at a final concentration of 50 ng/µl and pre-amplified by PCR (forward primer: CAATCCGCCCTCACTACAACCG, reverse primer: CTACTCTGGCGTCGATGAGGGA). PCR products were purified by agarose gel electrophoresis and the Nucleospin Gel and PCR clean-up kit (Macherey Nagel) and subcloned into pcDNA3.1 for pseudotyped lentiviral particle production. For the cell-based binding assay HA molecules were fused by a 6xGS linker to one of the three fluorescent proteins mTagBFP2 (FPbase ID: ZO7NN), superfolder (sf)GFP (FPBase ID: B4SOW), or dTomato (FPBase ID: G1DQY). For recombinant protein expression, HA coding regions excluding the transmembrane and C-terminal domains were subcloned for A/Hawaii/70/2019 (aa 1-529), A/Hong Kong/4801/2014 (aa 1-530) and A/Texas/37/2024 (aa 1-530) into pCAGGS and fused C-terminally to a Foldon trimerization domain, a sfGFP, a HRV 3C site, 8x His Tag and a Twin-Strep-tag.

### Recombinant protein production

Recombinant HA proteins were produced by transfection of HEK293-6E cells at a concentration of 0.8*10^6^ cells/ml. For transfection, per 1 ml of cell suspension, 1 µg HA expression plasmid was mixed with 45 µl DPBS (Gibco) and 3.4 µl of 0.45 mg/ml polyethylenimine (PEI, Sigma-Aldrich). After an incubation for 10 min at room temperature (RT), the transfection mixture was added dropwise to the cells and cells were incubated at 37°C and 6% CO_2_ in a shaking incubator at 110 rpm for 7 days. The cell culture supernatant was harvested by centrifugation and filtered through a 0.45 µm PVDF membrane. HA proteins were purified on chromatography columns (BioRad) using Strep-Tactin®XT 4Flow® resin (IBA Lifesciences) according to the manufacturer’s instructions. After elution, proteins were concentrated in PBS using 30K Amicon centrifugal filters (Millipore) and their concentration was determined by UV spectrophotometry (Nanodrop, Thermo-Fisher). Recombinant SARS-CoV-2 S6P was kindly provided by Manuel Koch and has been produced and purified as previously described.^77^

### Purification of poly-IgGs from plasma and plasma-IgG depletion

For poly-IgG purification, plasma samples were diluted in PBS and incubated overnight at 4°C with Protein G Sepharose™ 4 Fast Flow (Cytiva) on a rotating platform. Samples were then loaded onto chromatography columns (BioRad) and the flow through fraction was collected as IgG-depleted plasma. After washing the Sepharose beads with DPBS (Gibco), poly-IgGs were eluted with 0.1 M glycine (pH 3) into 1 M Tris HCl (pH 8). Poly-IgG and IgG-depleted plasma samples were concentrated in PBS using 30K Amicon centrifugal filters (Millipore) and stored at 4°C. The concentration was measured by UV spectrophotometry (Nanodrop, Thermo-Fisher).

### Depletion of HA-reactive antibodies from poly-IgGs

To deplete poly-IgG samples from HA-reactive antibodies, 1 ml bed volume of Strep-Tactin^®^XT 4Flow^®^ resin (IBA Lifesciences) was loaded onto chromatography columns (BioRad). After washing with 1x Buffer W (100 mM Tris/HCl, 150 mM NaCl, 1mM EDTA), a mix of the three produced recombinant HA proteins (2 mg each of A/Hawaii/70/2019 and A/Hong Kong/4801/2014, 1 mg of A/Texas/37/2024) was added to the resin and incubated for 30 min. The flowthrough was discarded and 2 mg of poly-IgG sample diluted in 1x Buffer W was added to the column followed by 12 ml of 1x Buffer W. The flowthrough was collected and concentrated during buffer exchange to PBS using 30K Amicon centrifugal filters (Millipore). IgG concentrations were determined by UV spectrophotometry (Nanodrop, Thermo-Fisher).

### Enzyme-linked immunosorbent assay

For measuring binding reactivity, high binding 96-well plates (Corning) were coated overnight at 4°C with 2.5 µg/ml of protein (Influenza HA or SARS-CoV-2 S HexaPro protein). Plates were washed 3x with PBST (PBS with 0.2% Tween-20 (Carl Roth)) and blocked for 1 h with blocking buffer (PBST with 2.5% BSA (Sigma-Aldrich) and 2.5% dry milk powder (PanReac AppliChem)) at RT. After 3 washing steps with PBST, plates were incubated with serial dilutions of samples (i.e., plasma, purified poly-IgG or monoclonal antibodies) in blocking buffer for 2 h. For competitive ELISA experiments, primary antibodies were incubated for only 1h, washed 3 times with PBST, and incubated for 1 h with fixed concentrations of competitive monoclonal antibodies that have been biotinylated with the EZ-Link™ Sulfo-NHS-LC-Biotinylation Kit (Thermo Fisher Scientific). Plates were washed 3 times with PBST and human IgG samples were detected with 1:2000 goat anti-human IgG-HRP (Southern Biotech) in blocking buffer, while biotinylated antibodies were detected with 1:5000 Pierce™ High Sensitivity Streptavidin-HRP (Thermo Fisher Scientific) in PBST with 0.5% BSA (Sigma-Aldrich) and 0.5% dry milk powder (PanReac AppliChem). After 0.5 - 1 h incubation at RT, plates were washed three times with PBST, 150 µl ABTS substrate was added to each well, and absorbance at 415 nm with reference at 695 nm was measured on a microplate reader (Tecan). For each plate, a sandwich ELISA for a standardized human IgG (IgG1, Kappa from human myeloma plasma; Sigma-Aldrich) was performed in dilutions from 5 to 0.000064 µg/ml and plates were developed to a top OD_415-695_ of approximately 1.5 of the standard.

### Production and titration of HA-pseudotyped lentiviral particles

Lentivirus-based pseudovirus particles were produced by co-transfection of plasmids coding for HIV-1 Tat, HIV-1 Gag/Pol, HIV-1 Rev, luciferase followed by an IRES and ZsGreen as described previously,^78,79^ but adapted for the expression of HA protein of the respective influenza A virus strain and the addition of neuraminidase for viral release and trypsin for HA processing as described elsewhere.^80^ 2-4 h after a change to FreeStyle 293 Expression Medium (Thermo Fisher) HEK293T cells were transfected with the pseudovirus encoding plasmids using FuGENE 6 Transfection Reagent (Promega) and incubated at 37°C and 5% CO_2_. For HA-expressing viral particles, 25 mU neuraminidase from Clostridium perfringens (Sigma Aldrich) was added per 1 mL of cell culture medium at 16 h post transfection. 48 h post transfection, pseudovirus containing supernatant was harvested, filtered by a 0.45 µM PVDF syringe filter and HA-pseudotyped particles were incubated with 250 µg/ml TPCK-treated Trypsin (Sigma-Aldrich) at 37°C for 1 h. After stopping the trypsinization by incubation with 250 µg/ml of Soybean Trypsin Inhibitor (Roche) for 30 min at 37°C, the supernatant was stored at −80°C. By infecting MDCK-SIAT1 cells with serial dilutions of the virus supernatant, each virus batch was titrated. After an incubation for 48 h at 37°C and 5% CO_2_, cells were incubated for 2 min with luciferin/lysis buffer (10 mM MgCl_2_, 0.3 mM ATP, 0.5 mM Coenzyme A, 17 mM IGEPAL (all Sigma-Aldrich), and 1 mM D-Luciferin (GoldBio) in Tris-HCL) and luciferase activity was measured using a microplate reader (Tristar 5, Berthold Technologies, counting time: 1 s). For neutralization assay, viruses were diluted in DMEM (Gibco) with 1 mM L-Glutamine to yield a 1000-fold higher RLUs than that of uninfected cells, corresponding to approximately 50,000 - 100,000 RLUs.

### Pseudovirus neutralization assay

To test the neutralizing activity of serum and monoclonal antibodies in a microneutralization assay, serial dilutions of serum (heat-inactivated for 1 h at 56°C) and monoclonal antibodies were prepared in DMEM (Gibco) containing 1 mM L-Glutamine in a 96-well format. After an incubation with pseudovirus for 1 h at 37°C and 5% CO_2_, 1.25*10^4^ MDCK-SIAT1 cells were added per well and incubated at 37°C and 5% CO_2_ for another 48 h. SARS-CoV-2 assays were performed with ACE2-expressing HEK293T-cells.^78,79^ Luciferase activity was measured as described above. On each assay plate, negative controls (cells only and virus only) and virus positive controls (infected cells without serum/monoclonal antibody) were measured. The mean RLU of negative controls was subtracted as background signal and 50% inhibitory dilution (ID_50_) and 50% inhibitory concentration (IC_50_) values were calculated as the serum dilution or monoclonal antibody concentration leading to a 50% reduction of RLUs in comparison to the virus positive control.

### Virus neutralization test

Virus neutralizing antibodies in human serum samples and purified plasma-IgG monoclonal antibodies were investigated by a virus neutralization test on MDCK II cells (Collection of Cell Lines in Veterinary Medicine CCLV-RIE 1061). Human serum samples were serially diluted 2-fold in DMEM (starting concentration 1:16), and purified plasma-IgG monoclonal antibodies were adjusted to a concentration of 100 µg/ml per sample, following serial dilution. Samples were analyzed in triplicates. Each diluted sample was mixed with 100 TCID_50_ of A/dairy cow/Texas/063224-24-1/2024 genotype B3.13 (H5N1, GISAID-No: EPI_ISL_19155861), the European HPAIV isolate A/wild goose/Germany-NW/2024AI02730/2024 genotype DE23-11-N1.3 euDG (H5N1, GISAID-No: EPI_ISL_19494411), or mixed with 100 TCID_50_ of A/Sydney/5/2021 (H1N1, GISAID-No: EPI_ISL_6424984), followed by an incubation period of 1 h at 37 °C. Subsequently, 100 µL of MDCK II cells were added per well, followed by a second incubation period of 72 h at 37 °C. Neutralizing antibodies were investigated by light microscopy and considered to be neutralizing in the absence of a cytopathic effect (CPE). The corresponding complete (100%) inhibitory dilution (ID_100_; reciprocal serum dilution) was defined as the dilution achieving 100% neutralization and was determined from the last serum dilution in which no CPE was observed. Negative and positive control sera were included and the virus titer was confirmed by titration.

### Cell-based binding assay

To test the binding activity of plasma and monoclonal antibodies, HEK293E cells were transfected with plasmids encoding for the desired HA-fluorophore fusion protein (coupled to either mTagBFP2, sfGFP, or dTomato) using 3.4 µl of 0.45 mg/ml PEI (Sigma-Aldrich) as transfection reagent and 0.5 µg of plasmid per 1 ml of cell suspension. After 48 h of incubation the cells were harvested by centrifugation. 600.000 cells of each HA and fluorescent protein were seeded together in one well in FACS buffer (PBS, 2 mM EDTA, 2% FCS). After centrifugation and removal of the supernatant serial dilutions of plasma with a starting dilution of 1:20 or monoclonal antibodies with the starting concentration of 100 µg/ml in PBS were added to the cells and incubated for 30 min on ice. After two washing steps the detection antibody (IgG, clone G18-145, 1:300 in FACS buffer, BV711, BD) was added to the cells and incubated for 30 min on ice. After washing and removal of the supernatant the live-dead marker (Zombie NIR, 1:600 in PBS, BioLegend) was added to the cells and incubated for 30 min on ice. After washing and removal of the supernatant cells were fixed with Cytofix fixation buffer (BD) for 30 min on ice. After two washing steps the cells were resuspended in FACS buffer and filtered through a 40 µm multiscreen-mesh filter plate (Merck Millipore). Cells were acquired on a LSR Fortessa (BD) equipped with a HTS (BD) with a set stopping gate of 10,000 living cells.

### Isolation of H5N1 A/Texas/37/2024 HA-specific IgG^+^ B cells

B cells from donors were magnetically enriched from PBMCs using CD19-microbeads (Miltenyi Biotec) according to the manufacturer’s instructions. Enriched B cells were stained with a mix of 4’,6-Diamindin-2-phenylindol (DAPI, Miltenyi Biotec), CD20 (BD, clone 2H7, Alexa Fluor 700), IgG (BD, clone G18-145, PE) and DyLight488- and DyLight650-labeled H5N1 A/Texas/37/2024(Y98F) HA protein (10 µg/ml). DAPI^-^, CD20^+^, IgG^+^, and double positive H5N1 A/Texas/37/2024(Y98F) protein cells were single cell sorted in a 96-well plate using an Aria Fusion (Becton Dickinson). Wells of the 96-well plate contained 4 µl buffer with 0.5x PBS, 0.5 U/µl RNAsin (Promega), 0.5 U/µl RNaseOUT (Thermo Fisher Scientific), and 10 mM DTT (Thermo Fisher Scientific). Immediately after sorting, the plates were stored at −80°C until further processing.

### Single cell B cell receptor sequencing and cloning of monoclonal antibodies

B cell receptor sequencing and cloning of monoclonal antibodies has been performed as described in detail elsewhere.^37^ In brief, cDNA from single cells was generated with random hexamer primers (Thermo Fisher Scientific) and Superscript IV (Thermo Fisher Scientific). B cell receptor heavy and light chain sequences were amplified in a semi-nested PCR with Platinum Taq Polymerase (Thermo Fisher Scientific) using human V gene segment specific forward and IgG constant region reverse primers.^81^ PCR products were sequenced by Sanger sequencing and processed with the Antibody Analysis Toolkit (AbRAT)^82^ for quality control and sequence annotation (based on IgBLAST^83^ and BLAST^84^). Variable regions were either amplified from first PCR products with primers containing overhangs for sequence and ligation independent cloning (SLIC) as described previously^37^ or ordered as gene fragments without endogenous leader signal from IDT as eBlocks. For previously published antibodies, amino acid sequences of heavy and light chain variable regions were derived from the respective publication and/or public repositories (PDB.org; Supplementary Table 3), were reverse transcribed as well as codon optimized for human cell expression with the IDT codon optimization webtool,^85^ and ordered from IDT as eBlocks. PCR products with SLIC overhangs and ordered gene fragments were cloned into IgG1, IgK, or IgL antibody expression plasmids from Tiller et al.^86^ by SLIC using either the endogenous leader sequence (PCR products) or the plasmid backbone derived murine leader sequence (gene fragments) as described previously.^37,87^

### Production and purification of monoclonal antibodies

Monoclonal antibodies were produced by co-transfection of HEK293-6E cells at a concentration of 0.8*10^6^ cells/ml or HEK293T cells at 80% confluency with heavy and light chain expression plasmids using PEI (Sigma-Aldrich) as described previously.^37,87,88^ For HEK293-6E-cells, per 1 ml of cell suspension, 0.5 µg each of the heavy and light chain plasmid was mixed with 45 µl DPBS (Gibco). After adding 3.4 µl of 0.45 mg/ml PEI the transfection mixture was incubated for 10 min at RT and then added dropwise to the cells. The cells were kept at 37°C and 6% CO_2_ in a shaking incubator at 110 rpm for 7 days. For HEK293T cells, after washing cells with serum free DMEM for 1 h, the medium was replaced with DMEM supplemented with 1% Nutridoma-SP (Roche), 1 mM sodium pyruvate, 1% Penicillin-Streptomycin (Gibco) and 1 mM L-Glutamine (Gibco). Per 15 cm cell culture dish, 15 µg each of the heavy and light chain plasmid was mixed with 1.2 ml DPBS (Gibco). 200 µl of 0.45 mg/ml PEI was added and after vortexing for 20 sec and an incubation for 10 min at RT the mix was then added dropwise to the cells. The cells were incubated at 37°C and 5% CO_2_ for 3 days.

Cell culture supernatants were harvested by centrifugation and incubated overnight at 4°C with Protein G Sepharose™ 4 Fast Flow (Cytiva) on a rotating platform. Antibodies were then purified using chromatography columns (Bio-Rad). After washing the Sepharose beads with DPBS (Gibco), antibodies were eluted with 0.1 M Glycine (pH 3) into 1 M Tris HCl (pH 8). Buffer exchange to PBS was performed using 30K Amicon centrifugal filters (Millipore). Antibodies were stored at 4°C and their concentration was measured by UV spectrophotometry (Nanodrop, Thermo-Fisher).

### In vivo challenge experiment

The animal experiment described here was evaluated by the responsible ethics committee of the State Office of Agriculture, Food Safety, and Fishery in Mecklenburg-Western Pomerania (LALLF M-V) and gained governmental approval under registration number LVL MV TSD/ 7221.3-1-042/24. 6-to-8-week-old female BALB/c mice (Charles River) were inoculated i.p. with 10 mg/kg monoclonal antibody (48H5-1-1G9K, 47H5-1-2B12K, N=4; or the SARS-CoV2-specific control antibody DZIF-10c^45,46^, N=5) in 100 µl physiological saline (0.9% sodium chloride) on day −1 and infected intranasally on d0 with 10^3^ TCID_50_ H5N1 (bovine isolate A/dairy cow/Texas/063224-24-1/2024, genotype B3.13). A summary of the animal numbers and the experimental design is given in Figure 6A. Animals were scored daily for weight loss and clinical symptoms. Euthanasia was performed when the animals lost 20% of their body weight or exhibited severe clinical signs. This was done under isoflurane anesthesia, followed by cervical dislocation. All remaining animals were euthanized on day 10 for terminal blood draw and lung tissue extraction.

### Quantification

Serum ED_50_ and poly-IgG/monoclonal antibody EC_50_ values from ELISA were calculated by a nonlinear sigmoidal four parameter logistic fit model with GraphPad Prism 10. Serum ED_50_/ID_50_ and poly-IgG/monoclonal antibody EC_50_/IC_50_ values from surface-expressed HA-binding and HA-pseudotyped lentivirus neutralization assays were determined using a normalized nonlinear fit model (agonist versus normalized response curve with variable slope) in GraphPad Prism 10. In cases where the calculated values were below the detection threshold, they were inferred by extrapolation from a partial dose response curve. Standard statistical errors for the titers of each individual were obtained as follows. From the data of Figure 1C, growth curves were resampled, assuming normally distributed measurement errors with standard deviations of 0.5 (log dilution titers) and 5% (growth rates), to learn the posterior distributions of individual EC_50_ values. The standard deviation was found to be related to the mean: Titers obtained by extrapolation from a partial dose response curve have a larger error margin than titers fitted from a full dose response curve. This error margin increases exponentially as the inferred EC_50_ deviates from the interval of experimentally measured concentrations. In all other panels in Figure 1, we used the error model learned from the data of Figure 1C to resample ED_50_/ID_50_ or EC_50_/IC_50_ values. Next, we obtained cohort population distributions of characteristic titers by aggregating the posterior EC_50_ distributions of the individuals in each cohort (n = 66 in the upper panel, n = 20 in the lower panel of Figure 1). Based on these distributions, we report the characteristic titer range in a given cohort in the form 50% percentile [25% percentile, 75% percentile].

## Acknowledgments

We thank Mareen Grawe, Lisa Kottege, Hanna Janicki and Banu Meiners for excellent technical assistance, Timm Harder for kindly providing the virus isolate A/Sydney/5/2021 (H1N1) and Manuel Koch for kindly providing recombinant SARS-CoV-2 S6P protein. We are grateful to Diego Diel for kindly providing virus isolate A/dairy cow/Texas/063224-24-1/2024 (H5N1).

## Author contributions

Conceptualization: CK, FK

Methodology: KD, LU, DR, MM, JE, ALS, MW, MG, ML, CK

Investigation: KD, LU, DR, MM, NH, UW, JE, JS, RS, MS, MW, LGL, EMA, MK, CD, AP, MR, LG, HG, CK

Formal Analysis: KD, LU, DR, MM, NH, JE, CK

Visualization: KD, LU, DR, CK

Funding acquisition: CK, MB, ML, FK

Project administration: HG, LG, CK, FK

Resources: DH, MB, TE, ML, CKu, FK Supervision: CKu, DH, MB, TE, ML, CK, FK

Writing – original draft: KD, LU, CK

Writing – review & editing: all authors

## Data and materials availability

Amino acid sequences of HA-variants used in this study are available from gisaid.org. All data supporting the findings of this study are included in the main text or supplementary materials. Requests for any additional material will be fulfilled by the corresponding authors upon request but might be restricted by sample availability, data protection, ethics vote, and require a Material Transfer Agreement (MTA) for non-commercial usage.

## References and Notes

1. Cumulative number of confirmed human cases for avian influenza A(H5N1) reported to WHO, 2003-2024, 12 December 2024 https://www.who.int/publications/m/item/cumulative-number-of-confirmed-human-cases-for-avian-influenza-a(h5n1)-reported-to-who--2003-2024--20-december-2024.

2. Uyeki, T.M., Milton, S., Abdul Hamid, C., Reinoso Webb, C., Presley, S.M., Shetty, V., Rollo, S.N., Martinez, D.L., Rai, S., Gonzales, E.R., et al. (2024). Highly Pathogenic Avian Influenza A(H5N1) Virus Infection in a Dairy Farm Worker. N Engl J Med, NEJMc2405371. 10.1056/NEJMc2405371.

3. Caliendo, V., Lewis, N.S., Pohlmann, A., Baillie, S.R., Banyard, A.C., Beer, M., Brown, I.H., Fouchier, R.A.M., Hansen, R.D.E., Lameris, T.K., et al. (2022). Transatlantic spread of highly pathogenic avian influenza H5N1 by wild birds from Europe to North America in 2021. Sci Rep 12, 11729. 10.1038/s41598-022-13447-z.

4. Nguyen, T.-Q., Hutter, C., Markin, A., Thomas, M., Lantz, K., Killian, M.L., Janzen, G.M., Vijendran, S., Wagle, S., Inderski, B., et al. (2024). Emergence and interstate spread of highly pathogenic avian influenza A(H5N1) in dairy cattle. Preprint, 10.1101/2024.05.01.591751 https://doi.org/10.1101/2024.05.01.591751.

5. CDC (2024). H5 Bird Flu: Current Situation. Avian Influenza (Bird Flu). https://www.cdc.gov/bird-flu/situation-summary/index.html.

6. Garg, S., Reinhart, K., Couture, A., Kniss, K., Davis, C.T., Kirby, M.K., Murray, E.L., Zhu, S., Kraushaar, V., Wadford, D.A., et al. (2024). Highly Pathogenic Avian Influenza A(H5N1) Virus Infections in Humans. N Engl J Med, NEJMoa2414610. 10.1056/NEJMoa2414610.

7. Mellis, A.M., Coyle, J., Marshall, K.E., Frutos, A.M., Singleton, J., Drehoff, C., Merced-Morales, A., Pagano, H.P., Alade, R.O., White, E.B., et al. (2024). Serologic Evidence of Recent Infection with Highly Pathogenic Avian Influenza A(H5) Virus Among Dairy Workers — Michigan and Colorado, June–August 2024. MMWR Morb. Mortal. Wkly. Rep. 73, 1004–1009. 10.15585/mmwr.mm7344a3.

8. Peacock, T., Moncla, L., Dudas, G., VanInsberghe, D., Sukhova, K., Lloyd-Smith, J.O., Worobey, M., Lowen, A.C., and Nelson, M.I. (2024). The global H5N1 influenza panzootic in mammals. Nature. 10.1038/s41586-024-08054-z.

9. Krammer, F. (2019). The human antibody response to influenza A virus infection and vaccination. Nat Rev Immunol 19, 383–397. 10.1038/s41577-019-0143-6.

10. Arzey, G.G., Kirkland, P.D., Arzey, K.E., Frost, M., Maywood, P., Conaty, S., Hurt, A.C., Deng, Y.-M., Iannello, P., Barr, I., et al. (2012). Influenza Virus A (H10N7) in Chickens and Poultry Abattoir Workers, Australia. Emerg. Infect. Dis. 18, 814–816. 10.3201/eid1805.111852.

11. Wei, S.-H., Yang, J.-R., Wu, H.-S., Chang, M.-C., Lin, J.-S., Lin, C.-Y., Liu, Y.-L., Lo, Y.-C., Yang, C.-H., Chuang, J.-H., et al. (2013). Human infection with avian influenza A H6N1 virus: an epidemiological analysis. The Lancet Respiratory Medicine 1, 771–778. 10.1016/S2213-2600(13)70221-2.

12. Hobson, D., Curry, R.L., Beare, A.S., and Ward-Gardner, A. (1972). The role of serum haemagglutination-inhibiting antibody in protection against challenge infection with influenza A2 and B viruses. Epidemiol. Infect. 70, 767–777. 10.1017/S0022172400022610.

13. Ohmit, S.E., Petrie, J.G., Cross, R.T., Johnson, E., and Monto, A.S. (2011). Influenza Hemagglutination-Inhibition Antibody Titer as a Correlate of Vaccine-Induced Protection. The Journal of Infectious Diseases 204, 1879–1885. 10.1093/infdis/jir661.

14. Fox, A., Mai, L.Q., Thanh, L.T., Wolbers, M., Le Khanh Hang, N., Thai, P.Q., Thu Yen, N.T., Minh Hoa, L.N., Bryant, J.E., Duong, T.N., et al. (2015). Hemagglutination inhibiting antibodies and protection against seasonal and pandemic influenza infection. Journal of Infection 70, 187–196. 10.1016/j.jinf.2014.09.003.

15. Nakamura, G., Chai, N., Park, S., Chiang, N., Lin, Z., Chiu, H., Fong, R., Yan, D., Kim, J., Zhang, J., et al. (2013). An In Vivo Human-Plasmablast Enrichment Technique Allows Rapid Identification of Therapeutic Influenza A Antibodies. Cell Host & Microbe 14, 93–103. 10.1016/j.chom.2013.06.004.

16. Dreyfus, C., Laursen, N.S., Kwaks, T., Zuijdgeest, D., Khayat, R., Ekiert, D.C., Lee, J.H., Metlagel, Z., Bujny, M.V., Jongeneelen, M., et al. (2012). Highly Conserved Protective Epitopes on Influenza B Viruses. Science 337, 1343–1348. 10.1126/science.1222908.

17. Throsby, M., Van Den Brink, E., Jongeneelen, M., Poon, L.L.M., Alard, P., Cornelissen, L., Bakker, A., Cox, F., Van Deventer, E., Guan, Y., et al. (2008). Heterosubtypic Neutralizing Monoclonal Antibodies Cross-Protective against H5N1 and H1N1 Recovered from Human IgM+ Memory B Cells. PLoS ONE 3, e3942. 10.1371/journal.pone.0003942.

18. Ohshima, N., Iba, Y., Kubota-Koketsu, R., Asano, Y., Okuno, Y., and Kurosawa, Y. (2011). Naturally Occurring Antibodies in Humans Can Neutralize a Variety of Influenza Virus Strains, Including H3, H1, H2, and H5. J Virol 85, 11048–11057. 10.1128/JVI.05397-11.

19. Corti, D., Suguitan, A.L., Pinna, D., Silacci, C., Fernandez-Rodriguez, B.M., Vanzetta, F., Santos, C., Luke, C.J., Torres-Velez, F.J., Temperton, N.J., et al. (2010). Heterosubtypic neutralizing antibodies are produced by individuals immunized with a seasonal influenza vaccine. J. Clin. Invest. 120, 1663–1673. 10.1172/JCI41902.

20. Corti, D., Voss, J., Gamblin, S.J., Codoni, G., Macagno, A., Jarrossay, D., Vachieri, S.G., Pinna, D., Minola, A., Vanzetta, F., et al. (2011). A Neutralizing Antibody Selected from Plasma Cells That Binds to Group 1 and Group 2 Influenza A Hemagglutinins. Science 333, 850–856. 10.1126/science.1205669.

21. Fu, Y., Zhang, Z., Sheehan, J., Avnir, Y., Ridenour, C., Sachnik, T., Sun, J., Hossain, M.J., Chen, L.-M., Zhu, Q., et al. (2016). A broadly neutralizing anti-influenza antibody reveals ongoing capacity of haemagglutinin-specific memory B cells to evolve. Nat Commun 7, 12780. 10.1038/ncomms12780.

22. Hu, W., Chen, A., Miao, Y., Xia, S., Ling, Z., Xu, K., Wang, T., Xu, Y., Cui, J., Wu, H., et al. (2013). Fully human broadly neutralizing monoclonal antibodies against influenza A viruses generated from the memory B cells of a 2009 pandemic H1N1 influenza vaccine recipient. Virology 435, 320–328. 10.1016/j.virol.2012.09.034.

23. Garcia, J.-M., Pepin, S., Lagarde, N., Ma, E.S.K., Vogel, F.R., Chan, K.H., Chiu, S.S.S., and Peiris, J.S.M. (2009). Heterosubtype Neutralizing Responses to Influenza A (H5N1) Viruses Are Mediated by Antibodies to Virus Haemagglutinin. PLoS ONE 4, e7918. 10.1371/journal.pone.0007918.

24. Nefkens, I., Garcia, J.-M., Ling, C.S., Lagarde, N., Nicholls, J., Tang, D.J., Peiris, M., Buchy, P., and Altmeyer, R. (2007). Hemagglutinin pseudotyped lentiviral particles: Characterization of a new method for avian H5N1 influenza sero-diagnosis. Journal of Clinical Virology 39, 27–33. 10.1016/j.jcv.2007.02.005.

25. Sui, J., Sheehan, J., Hwang, W.C., Bankston, L.A., Burchett, S.K., Huang, C.-Y., Liddington, R.C., Beigel, J.H., and Marasco, W.A. (2011). Wide Prevalence of Heterosubtypic Broadly Neutralizing Human Anti-Influenza A Antibodies. Clinical Infectious Diseases 52, 1003–1009. 10.1093/cid/cir121.

26. Daulagala, P., Cheng, S.M.S., Chin, A., Luk, L.L.H., Leung, K., Wu, J.T., Poon, L.L.M., Peiris, M., and Yen, H.-L. Avian Influenza A(H5N1) Neuraminidase Inhibition Antibodies in Healthy Adults after Exposure to Influenza A(H1N1)pdm09 - Volume 30, Number 1—January 2024 - Emerging Infectious Diseases journal - CDC. 10.3201/eid3001.230756.

27. Zhang, R., Rong, X., Pan, W., and Peng, T. (2011). Determination of serum neutralization antibodies against seasonal influenza A strain H3N2 and the emerging strains 2009 H1N1 and avian H5N1. Scandinavian Journal of Infectious Diseases 43, 216–220. 10.3109/00365548.2010.539258.

28. Treanor, J.J., Campbell, J.D., Zangwill, K.M., Rowe, T., and Wolff, M. (2006). Safety and Immunogenicity of an Inactivated Subvirion Influenza A (H5N1) Vaccine. N Engl J Med 354, 1343–1351. 10.1056/NEJMoa055778.

29. Bresson, J.-L., Perronne, C., Launay, O., Gerdil, C., Saville, M., Wood, J., Höschler, K., and Zambon, M.C. (2006). Safety and immunogenicity of an inactivated split-virion influenza A/Vietnam/1194/2004 (H5N1) vaccine: phase I randomised trial. The Lancet 367, 1657–1664. 10.1016/S0140-6736(06)68656-X.

30. CDC (2024). CDC A(H5N1) Bird Flu Response Update June 14, 2024. Avian Influenza (Bird Flu). https://www.cdc.gov/bird-flu/spotlights/h5n1-response-06142024.html.

31. Sanz-Muñoz, I., Sánchez-Martínez, J., Rodríguez-Crespo, C., Concha-Santos, C.S., Hernández, M., Rojo-Rello, S., Domínguez-Gil, M., Mostafa, A., Martinez-Sobrido, L., Eiros, J.M., et al. (2024). Are we serologically prepared against an avian influenza pandemic and could seasonal flu vaccines help us? mBio, e03721–24. 10.1128/mbio.03721-24.

32. Matrosovich, M., Matrosovich, T., Carr, J., Roberts, N.A., and Klenk, H.-D. (2003). Overexpression of the α-2,6-Sialyltransferase in MDCK Cells Increases Influenza Virus Sensitivity to Neuraminidase Inhibitors. Journal of Virology 77, 8418–8425. 10.1128/jvi.77.15.8418-8425.2003.

33. Halwe, N.J., Cool, K., Breithaupt, A., Schön, J., Trujillo, J.D., Nooruzzaman, M., Kwon, T., Ahrens, A.K., Britzke, T., McDowell, C.D., et al. (2024). H5N1 clade 2.3.4.4b dynamics in experimentally infected calves and cows. Nature, 1–3. 10.1038/s41586-024-08063-y.

34. Whittle, J.R.R., Wheatley, A.K., Wu, L., Lingwood, D., Kanekiyo, M., Ma, S.S., Narpala, S.R., Yassine, H.M., Frank, G.M., Yewdell, J.W., et al. (2014). Flow cytometry reveals that H5N1 vaccination elicits cross-reactive stem-directed antibodies from multiple Ig heavy-chain lineages. J Virol 88, 4047–4057. 10.1128/JVI.03422-13.

35. Wheatley, A.K., Whittle, J.R.R., Lingwood, D., Kanekiyo, M., Yassine, H.M., Ma, S.S., Narpala, S.R., Prabhakaran, M.S., Matus-Nicodemos, R.A., Bailer, R.T., et al. (2015). H5N1 Vaccine–Elicited Memory B Cells Are Genetically Constrained by the IGHV Locus in the Recognition of a Neutralizing Epitope in the Hemagglutinin Stem. The Journal of Immunology 195, 602–610. 10.4049/jimmunol.1402835.

36. Martín, J., Wharton, S.A., Lin, Y.P., Takemoto, D.K., Skehel, J.J., Wiley, D.C., and Steinhauer, D.A. (1998). Studies of the binding properties of influenza hemagglutinin receptor-site mutants. Virology 241, 101–111. 10.1006/viro.1997.8958.

37. Gieselmann, L., Kreer, C., Ercanoglu, M.S., Lehnen, N., Zehner, M., Schommers, P., Potthoff, J., Gruell, H., and Klein, F. (2021). Effective high-throughput isolation of fully human antibodies targeting infectious pathogens. Nat Protoc 16, 3639–3671. 10.1038/s41596-021-00554-w.

38. Hu, H., Voss, J., Zhang, G., Buchy, P., Zuo, T., Wang, L., Wang, F., Zhou, F., Wang, G., Tsai, C., et al. (2012). A Human Antibody Recognizing a Conserved Epitope of H5 Hemagglutinin Broadly Neutralizes Highly Pathogenic Avian Influenza H5N1 Viruses. J Virol 86, 2978–2989. 10.1128/JVI.06665-11.

39. Kallewaard, N.L., Corti, D., Collins, P.J., Neu, U., McAuliffe, J.M., Benjamin, E., Wachter-Rosati, L., Palmer-Hill, F.J., Yuan, A.Q., Walker, P.A., et al. (2016). Structure and Function Analysis of an Antibody Recognizing All Influenza A Subtypes. Cell 166, 596–608. 10.1016/j.cell.2016.05.073.

40. Thornburg, N.J., Nannemann, D.P., Blum, D.L., Belser, J.A., Tumpey, T.M., Deshpande, S., Fritz, G.A., Sapparapu, G., Krause, J.C., Lee, J.H., et al. (2013). Human antibodies that neutralize respiratory droplet transmissible H5N1 influenza viruses. J. Clin. Invest. 123, 4405– 4409. 10.1172/JCI69377.

41. Krause, J.C., Tsibane, T., Tumpey, T.M., Huffman, C.J., Basler, C.F., and Crowe, J.E. (2011). A Broadly Neutralizing Human Monoclonal Antibody That Recognizes a Conserved, Novel Epitope on the Globular Head of the Influenza H1N1 Virus Hemagglutinin. J Virol 85, 10905–10908. 10.1128/JVI.00700-11.

42. Ekiert, D.C., Friesen, R.H.E., Bhabha, G., Kwaks, T., Jongeneelen, M., Yu, W., Ophorst, C., Cox, F., Korse, H.J.W.M., Brandenburg, B., et al. (2011). A Highly Conserved Neutralizing Epitope on Group 2 Influenza A Viruses. Science 333, 843–850. 10.1126/science.1204839.

43. Bangaru, S., Lang, S., Schotsaert, M., Vanderven, H.A., Zhu, X., Kose, N., Bombardi, R., Finn, J.A., Kent, S.J., Gilchuk, P., et al. (2019). A Site of Vulnerability on the Influenza Virus Hemagglutinin Head Domain Trimer Interface. Cell 177, 1136–1152.e18. 10.1016/j.cell.2019.04.011.

44. Aartse, A., Eggink, D., Claireaux, M., van Leeuwen, S., Mooij, P., Bogers, W.M., Sanders, R.W., Koopman, G., and van Gils, M.J. (2021). Influenza A Virus Hemagglutinin Trimer, Head and Stem Proteins Identify and Quantify Different Hemagglutinin-Specific B Cell Subsets in Humans. Vaccines (Basel) 9, 717. 10.3390/vaccines9070717.

45. Kreer, C., Zehner, M., Weber, T., Ercanoglu, M.S., Gieselmann, L., Rohde, C., Halwe, S., Korenkov, M., Schommers, P., Vanshylla, K., et al. (2020). Longitudinal Isolation of Potent Near-Germline SARS-CoV-2-Neutralizing Antibodies from COVID-19 Patients. Cell 182, 843–854.e12. 10.1016/j.cell.2020.06.044.

46. Halwe, S., Kupke, A., Vanshylla, K., Liberta, F., Gruell, H., Zehner, M., Rohde, C., Krähling, V., Gellhorn Serra, M., Kreer, C., et al. (2021). Intranasal Administration of a Monoclonal Neutralizing Antibody Protects Mice against SARS-CoV-2 Infection. Viruses 13, 1498. 10.3390/v13081498.

47. De Jong, J.C., Claas, E.C.J., Osterhaus, A.D.M.E., Webster, R.G., and Lim, W.L. (1997). A pandemic warning? Nature 389, 554–554. 10.1038/39218.

48. Nachbagauer, R., Choi, A., Hirsh, A., Margine, I., Iida, S., Barrera, A., Ferres, M., Albrecht, R.A., García-Sastre, A., Bouvier, N.M., et al. (2017). Defining the antibody cross-reactome directed against the influenza virus surface glycoproteins. Nat Immunol 18, 464–473. 10.1038/ni.3684.

49. Yassine, H.M., McTamney, P.M., Boyington, J.C., Ruckwardt, T.J., Crank, M.C., Smatti, M.K., Ledgerwood, J.E., and Graham, B.S. (2018). Use of Hemagglutinin Stem Probes Demonstrate Prevalence of Broadly Reactive Group 1 Influenza Antibodies in Human Sera. Sci Rep 8, 8628. 10.1038/s41598-018-26538-7.

50. Andrews, S.F., Huang, Y., Kaur, K., Popova, L.I., Ho, I.Y., Pauli, N.T., Dunand, C.J.H., Taylor, W.M., Lim, S., Huang, M., et al. (2015). Immune history profoundly affects broadly protective B cell responses to influenza. Sci. Transl. Med. 7. 10.1126/scitranslmed.aad0522.

51. Kok, A., Wilks, S.H., Tureli, S., James, S.L., Bestebroer, T.M., Burke, D.F., Funk, M., van der Vliet, S., Spronken, M.I., Rijnink, W.F., et al. (2024). A vaccine antigen central in influenza A(H5) virus antigenic space confers subtype-wide immunity. bioRxiv, 2024.08.06.606696. 10.1101/2024.08.06.606696.

52. Luczo, J.M., and Spackman, E. (2024). Epitopes in the HA and NA of H5 and H7 avian influenza viruses that are important for antigenic drift. FEMS Microbiology Reviews 48, fuae014. 10.1093/femsre/fuae014.

53. Zost, S.J., Gilchuk, P., Case, J.B., Binshtein, E., Chen, R.E., Nkolola, J.P., Schäfer, A., Reidy, J.X., Trivette, A., Nargi, R.S., et al. (2020). Potently neutralizing and protective human antibodies against SARS-CoV-2. Nature 584, 443–449. 10.1038/s41586-020-2548-6.

54. Alberini, I., Del Tordello, E., Fasolo, A., Temperton, N.J., Galli, G., Gentile, C., Montomoli, E., Hilbert, A.K., Banzhoff, A., Del Giudice, G., et al. (2009). Pseudoparticle neutralization is a reliable assay to measure immunity and cross-reactivity to H5N1 influenza viruses. Vaccine 27, 5998–6003. 10.1016/j.vaccine.2009.07.079.

55. Pappas, L., Foglierini, M., Piccoli, L., Kallewaard, N.L., Turrini, F., Silacci, C., Fernandez-Rodriguez, B., Agatic, G., Giacchetto-Sasselli, I., Pellicciotta, G., et al. (2014). Rapid development of broadly influenza neutralizing antibodies through redundant mutations. Nature 516, 418–422. 10.1038/nature13764.

56. Ellebedy, A.H., Krammer, F., Li, G.-M., Miller, M.S., Chiu, C., Wrammert, J., Chang, C.Y., Davis, C.W., McCausland, M., Elbein, R., et al. (2014). Induction of broadly cross-reactive antibody responses to the influenza HA stem region following H5N1 vaccination in humans. Proc. Natl. Acad. Sci. U.S.A. 111, 13133–13138. 10.1073/pnas.1414070111.

57. Nachbagauer, R., Choi, A., Izikson, R., Cox, M.M., Palese, P., and Krammer, F. (2016). Age Dependence and Isotype Specificity of Influenza Virus Hemagglutinin Stalk-Reactive Antibodies in Humans. mBio 7, e01996–15. 10.1128/mBio.01996-15.

58. Sidney, J., Kim, A.-R., de Vries, R.D., Peters, B., Meade, P.S., Krammer, F., Grifoni, A., and Sette, A. (2024). Targets of influenza human T-cell response are mostly conserved in H5N1. mBio, e0347924. 10.1128/mbio.03479-24.

59. Khurana, S., King, L.R., Manischewitz, J., Posadas, O., Mishra, A.K., Liu, D., Beigel, J.H., Rappuoli, R., Tsang, J.S., and Golding, H. (2024). Licensed H5N1 vaccines generate cross-neutralizing antibodies against highly pathogenic H5N1 clade 2.3.4.4b influenza virus. Nat Med. 10.1038/s41591-024-03189-y.

60. Sun, M., Lai, H., Na, F., Li, S., Qiu, X., Tian, J., Zhang, Z., and Ge, L. (2023). Monoclonal Antibody for the Prevention of Respiratory Syncytial Virus in Infants and Children: A Systematic Review and Network Meta-analysis. JAMA Netw Open 6, e230023. 10.1001/jamanetworkopen.2023.0023.

61. Mulangu, S., Dodd, L.E., Davey, R.T., Tshiani Mbaya, O., Proschan, M., Mukadi, D., Lusakibanza Manzo, M., Nzolo, D., Tshomba Oloma, A., Ibanda, A., et al. (2019). A Randomized, Controlled Trial of Ebola Virus Disease Therapeutics. N Engl J Med 381, 2293– 2303. 10.1056/NEJMoa1910993.

62. Akinosoglou, K., Rigopoulos, E.-A., Kaiafa, G., Daios, S., Karlafti, E., Ztriva, E., Polychronopoulos, G., Gogos, C., and Savopoulos, C. (2022). Tixagevimab/Cilgavimab in SARS-CoV-2 Prophylaxis and Therapy: A Comprehensive Review of Clinical Experience. Viruses 15, 118. 10.3390/v15010118.

63. Friesen, R.H.E., Lee, P.S., Stoop, E.J.M., Hoffman, R.M.B., Ekiert, D.C., Bhabha, G., Yu, W., Juraszek, J., Koudstaal, W., Jongeneelen, M., et al. (2014). A common solution to group 2 influenza virus neutralization. Proc Natl Acad Sci U S A 111, 445–450. 10.1073/pnas.1319058110.

64. Song, G., He, W., Callaghan, S., Anzanello, F., Huang, D., Ricketts, J., Torres, J.L., Beutler, N., Peng, L., Vargas, S., et al. (2021). Cross-reactive serum and memory B-cell responses to spike protein in SARS-CoV-2 and endemic coronavirus infection. Nat Commun 12, 2938. 10.1038/s41467-021-23074-3.

65. Ercanoglu, M.S., Gieselmann, L., Dähling, S., Poopalasingam, N., Detmer, S., Koch, M., Korenkov, M., Halwe, S., Klüver, M., Di Cristanziano, V., et al. (2022). No substantial preexisting B cell immunity against SARS-CoV-2 in healthy adults. iScience 25, 103951. 10.1016/j.isci.2022.103951.

66. Doud, M.B., Lee, J.M., and Bloom, J.D. (2018). How single mutations affect viral escape from broad and narrow antibodies to H1 influenza hemagglutinin. Nat Commun 9, 1386. 10.1038/s41467-018-03665-3.

67. Gruell, H., Vanshylla, K., Korenkov, M., Tober-Lau, P., Zehner, M., Münn, F., Janicki, H., Augustin, M., Schommers, P., Sander, L.E., et al. (2022). SARS-CoV-2 Omicron sublineages exhibit distinct antibody escape patterns. Cell Host & Microbe 30, 1231–1241.e6. 10.1016/j.chom.2022.07.002.

68. Kikawa, C., Cartwright-Acar, C.H., Stuart, J.B., Contreras, M., Levoir, L.M., Evans, M.J., Bloom, J.D., and Goo, L. (2023). The effect of single mutations in Zika virus envelope on escape from broadly neutralizing antibodies. J Virol 97, e0141423. 10.1128/jvi.01414-23.

69. Cao, Y., Wang, J., Jian, F., Xiao, T., Song, W., Yisimayi, A., Huang, W., Li, Q., Wang, P., An, R., et al. (2022). Omicron escapes the majority of existing SARS-CoV-2 neutralizing antibodies. Nature 602, 657–663. 10.1038/s41586-021-04385-3.

70. Oraby, A.K., Stojic, A., Elawar, F., Bilawchuk, L.M., McClelland, R.D., Erwin, K., Granoski, M.J., Griffiths, C.D., Frederick, J.D., Arutyunova, E., et al. (2025). A single amino acid mutation alters multiple neutralization epitopes in the respiratory syncytial virus fusion glycoprotein. npj Viruses 3, 1–14. 10.1038/s44298-025-00119-8.

71. Stevens, J., Blixt, O., Tumpey, T.M., Taubenberger, J.K., Paulson, J.C., and Wilson, I.A. (2006). Structure and Receptor Specificity of the Hemagglutinin from an H5N1 Influenza Virus. Science 312, 404–410. 10.1126/science.1124513.

72. Glaser, L., Stevens, J., Zamarin, D., Wilson, I.A., García-Sastre, A., Tumpey, T.M., Basler, C.F., Taubenberger, J.K., and Palese, P. (2005). A single amino acid substitution in 1918 influenza virus hemagglutinin changes receptor binding specificity. J Virol 79, 11533–11536. 10.1128/JVI.79.17.11533-11536.2005.

73. Stevens, J., Blixt, O., Glaser, L., Taubenberger, J.K., Palese, P., Paulson, J.C., and Wilson, I.A. (2006). Glycan microarray analysis of the hemagglutinins from modern and pandemic influenza viruses reveals different receptor specificities. J Mol Biol 355, 1143–1155. 10.1016/j.jmb.2005.11.002.

74. Rogers, G.N., and D’Souza, B.L. (1989). Receptor binding properties of human and animal H1 influenza virus isolates. Virology 173, 317–322. 10.1016/0042-6822(89)90249-3.

75. Meijers, M., Ruchnewitz, D., Eberhardt, J., Karmakar, M., Łuksza, M., and Lässig, M. (2024). Concepts and methods for predicting viral evolution. Preprint at arXiv, 10.48550/ARXIV.2403.12684 https://doi.org/10.48550/ARXIV.2403.12684.

76. Worldwide Influenza Centre, CRICK Institute (2025). Annual and interim reports. Crick. https://www.crick.ac.uk/research/platforms-and-facilities/worldwide-influenza-centre/annual-and-interim-reports.

77. Vanshylla, K., Fan, C., Wunsch, M., Poopalasingam, N., Meijers, M., Kreer, C., Kleipass, F., Ruchnewitz, D., Ercanoglu, M.S., Gruell, H., et al. (2022). Discovery of ultrapotent broadly neutralizing antibodies from SARS-CoV-2 elite neutralizers. Cell Host & Microbe 30, 69–82.e10. 10.1016/j.chom.2021.12.010.

78. Vanshylla, K., Di Cristanziano, V., Kleipass, F., Dewald, F., Schommers, P., Gieselmann, L., Gruell, H., Schlotz, M., Ercanoglu, M.S., Stumpf, R., et al. (2021). Kinetics and correlates of the neutralizing antibody response to SARS-CoV-2 infection in humans. Cell Host & Microbe 29, 917–929.e4. 10.1016/j.chom.2021.04.015.

79. Crawford, K.H.D., Eguia, R., Dingens, A.S., Loes, A.N., Malone, K.D., Wolf, C.R., Chu, H.Y., Tortorici, M.A., Veesler, D., Murphy, M., et al. (2020). Protocol and Reagents for Pseudotyping Lentiviral Particles with SARS-CoV-2 Spike Protein for Neutralization Assays. Viruses 12, 513. 10.3390/v12050513.

80. Carnell, G.W., Ferrara, F., Grehan, K., Thompson, C.P., and Temperton, N.J. (2015). Pseudotype-Based Neutralization Assays for Influenza: A Systematic Analysis. Front. Immunol. 6. 10.3389/fimmu.2015.00161.

81. Kreer, C., Döring, M., Lehnen, N., Ercanoglu, M.S., Gieselmann, L., Luca, D., Jain, K., Schommers, P., Pfeifer, N., and Klein, F. (2020). openPrimeR for multiplex amplification of highly diverse templates. J Immunol Methods 480, 112752. 10.1016/j.jim.2020.112752.

82. Kreer, C. (2025). AbRAT: The Antibody Repertoire Analysis Toolkit. Version v1.0.1 (Zenodo). 10.5281/ZENODO.15311638 https://doi.org/10.5281/ZENODO.15311638.

83. Ye, J., Ma, N., Madden, T.L., and Ostell, J.M. (2013). IgBLAST: an immunoglobulin variable domain sequence analysis tool. Nucleic Acids Res 41, W34–40. 10.1093/nar/gkt382.

84. Camacho, C., Coulouris, G., Avagyan, V., Ma, N., Papadopoulos, J., Bealer, K., and Madden, T.L. (2009). BLAST+: architecture and applications. BMC Bioinformatics 10, 421. 10.1186/1471-2105-10-421.

85. Integrated DNA Technologies (IDT) Codon Optimization Tool.

86. Tiller, T., Meffre, E., Yurasov, S., Tsuiji, M., Nussenzweig, M.C., and Wardemann, H. (2008). Efficient generation of monoclonal antibodies from single human B cells by single cell RT-PCR and expression vector cloning. Journal of Immunological Methods 329, 112–124. 10.1016/j.jim.2007.09.017.

87. Korenkov, M., Zehner, M., Cohen-Dvashi, H., Borenstein-Katz, A., Kottege, L., Janicki, H., Vanshylla, K., Weber, T., Gruell, H., Koch, M., et al. (2023). Somatic hypermutation introduces bystander mutations that prepare SARS-CoV-2 antibodies for emerging variants. Immunity 56, 2803–2815.e6. 10.1016/j.immuni.2023.11.004.

88. Mouquet, H., Klein, F., Scheid, J.F., Warncke, M., Pietzsch, J., Oliveira, T.Y.K., Velinzon, K., Seaman, M.S., and Nussenzweig, M.C. (2011). Memory B Cell Antibodies to HIV-1 gp140 Cloned from Individuals Infected with Clade A and B Viruses. PLoS ONE 6, e24078. 10.1371/journal.pone.0024078.

