## Supplementary Information for "Detection of pre-existing humoral immunity against influenza virus H5N1 clade 2.3.4.4b in unexposed individuals"

### Supplementary Figures


| | |
|---|---|
| **Figure S1** | Cloning and purification of recombinant H5N1 (A/Texas/37/2024), H1N1 (A/Hawaii/70/2019) and H3N2 (A/Hong Kong/4801/2014) hemagglutinin proteins, related to Figure 1. |
| **Figure S2** | Example gating strategy for HEK293-6E cell surface-expressed HA binding assay, related to Figure 1 |
| **Figure S3** | Impact of sex, vaccination status, and age on A/Texas/37/2024 neutralization, related to Figure 1 |
| **Figure S4** | Comparison of serum vs. plasma-derived poly-IgG neutralization, related to Figure 1 |
| **Figure S5** | HA-pseudotyped virus neutralization and HA IgG binding before and after IgG depletion from plasma, related to Figure 1 |
| **Figure S6** | HA-pseudotyped virus neutralization and HA binding before and after depletion of HA-reactive antibodies from poly-IgG, realted to Figure 1 |
| **Figure S7** | HA subtype neutralization per individual, related to Figure 3 |
| **Figure S8** | Correlation of $ID_{50}$ and $ID_{100}$ values from H1N1 pseudovirus and authentic virus neutralization assay, related to Figure 3 |
| **Figure S9** | Example gating strategy for sorting of H5N1 A/Texas/37/2024 HA(Y98F)-specific memory B cells, related to Figure 4 |
| **Figure S10** | H5-reactive and clinically developed monoclonal antibodies bind and neutralize A/Texas/37/2024 HA, related to Figure 5 |
| **Figure S11** | H5N1 (euDG) and H1N1 authentic virus as well as H1N1 HA pseudotyped virus neutralization by monoclonal antibodies, related to Figure 5 |


### Supplementary Tables


| | |
|---|---|
| **Table S1** | Blood donor information |
| **Table S2** | Influenza strain information |
| **Table S3** | Monoclonal reference antibodies |

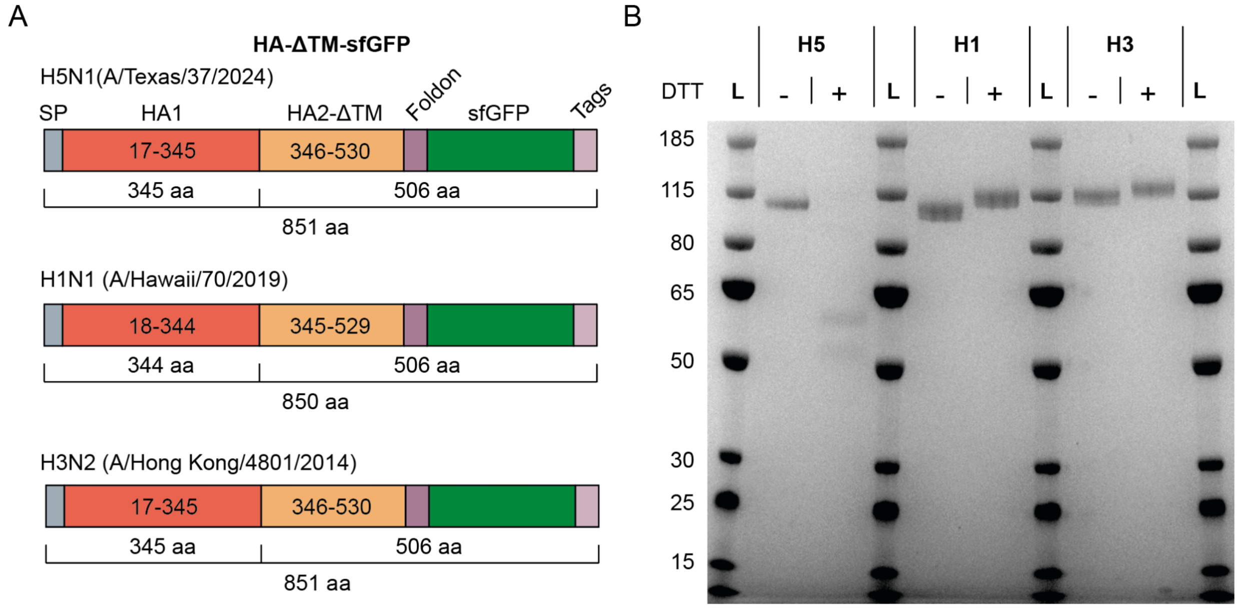


**Figure S1. Cloning and purification of recombinant H5N1 (A/Texas/37/2024), H1N1 (A/Hawaii/70/2019) and H3N2 (A/Hong Kong/4801/2014) hemagglutinin proteins, related to Figure 1.** (**A**) Cloned hemagglutinin (HA) proteins comprise the original HA coding sequences including the endogenous signal peptide (SP) but excluding the transmembrane and intracellular domain (ΔTM), i.e., amino acids (aa) 1-530 for H5, aa 1-529 for H1 and aa 1-530 for H3. Proteins were fused to a Foldon sequence for trimerization, a super-folder (sf)GFP, and C-terminal tags comprising an HRV 3C site, an 8xHis-Tag and a twin Strep-Tag. (**B**) Proteins were expressed in HEK293-6E suspension cells and purified on streptactin resin through the twin Strep-Tag. Purified proteins were analyzed by SDS-PAGE with or without reducing agent (dithiothreitol, DTT). Note that H5N1 contains a polybasic cleavage site. Hence, HA1 and HA2 are detected upon reduction, while the full-length HA0 is detected for H1 and H3. L = protein ladder.

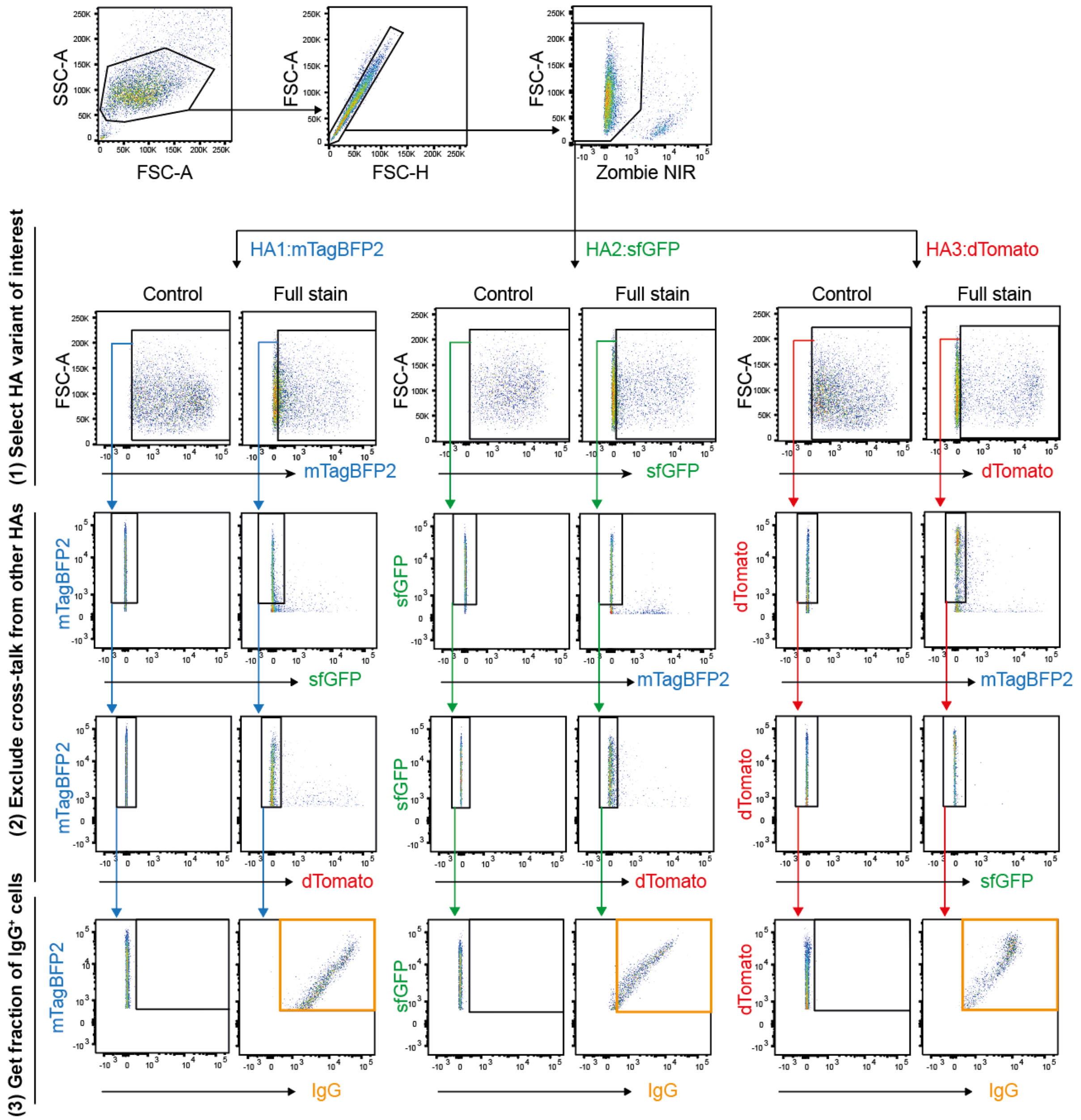


**Figure S2. Example gating strategy for the HEK293-6E cell surface-expressed HA binding assay, related to Figure 1.** Gating for IgG binding to cell surface-expressed HA starts with the isolation of the main HEK293-6E cell population by forward (FSC-A) and side scatter area (SSC-A), exclusion of doublets through forward scatter height (FSC-H) versus area (FSC-A) discrimination, and the exclusion of Zombie NIR positive (i.e., dead) cells. To identify the HA-expressing cells of interest, cells are selected for (1) the expression of the respective conjugated fluorescent protein (FP) using non-multiplexed cells expressing one of the other fluorophores as the negative control (not shown) and (2) the absence of any signal from the other multiplexed HA-FP conjugates. By gating on the fluorescently labeled anti-IgG antibody (3), the fraction of IgG-bound cells was determined. Multiplexed cells without any IgG or serum dilutions were used as control to assess the background staining of the anti-IgG antibody.

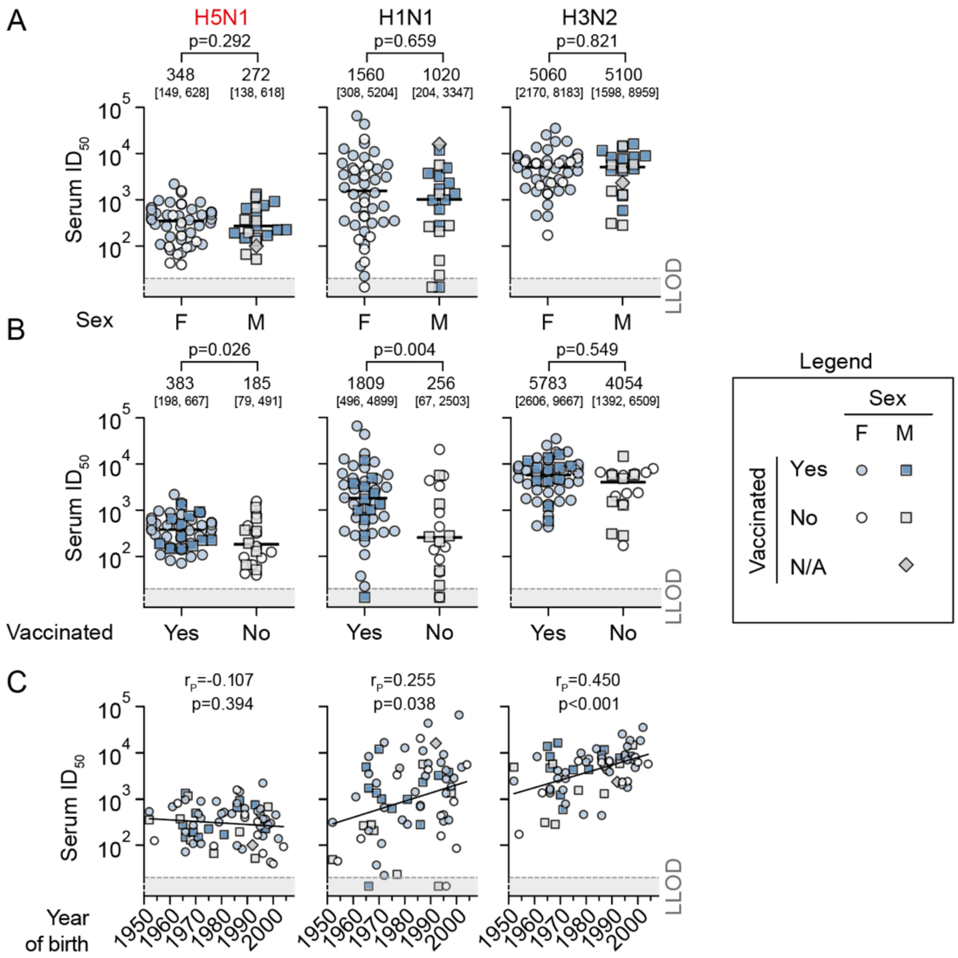


**Figure S3. Impact of sex, vaccination status, and age on A/Texas/37/2024 neutralization, related to Figure 1.** (**A**) Serum ID$_{50}$ values for A/Texas/37/2024 (H5N1), A/Hawaii/70/2019 (H1N1), and A/Hong Kong/4801/2014 (H3N2) HA-pseudotyped lentivirus, separated by the self-reported sex (female, n = 45; male n = 21). (**B**) ID$_{50}$ values as in (A), separated by self-reported vaccination status (no, n = 19; yes, n = 46). One individual did not report any information on vaccination and was excluded from the analysis in (B). Note that the last vaccination/exposure date varies between individuals (see also Table S1). P values in (A) and (B) represent the probabilities of observing median differences at least as extreme as those shown, assuming both samples come from the same distribution. These values were calculated using a two-sided Fisher's permutation test with 1,000 permutations. Horizontal bars in (A) and (B) represent the error-corrected median (see methods section for details), which is also reported on top of the graphs together with the $25^{th}$ and $75^{th}$ percentiles in brackets. (C) Correlation of log$_{10}$(ID$_{50}$) values from all n = 66 individuals with the year of birth. The line represents a linear regression of log$_{10}$ transformed ID$_{50}$ values. Pearson correlation coefficients $r_p$ and corresponding p-values of the transformed data are reported in the figures. The lowest tested dilution (considered as the lower limit of detection, LLOD) was a 20-fold dilution (dashed line in the plots). Calculated $ID_{50}$ values below the LLOD were arbitrarily set to a constant value below the LLOD for visualization.

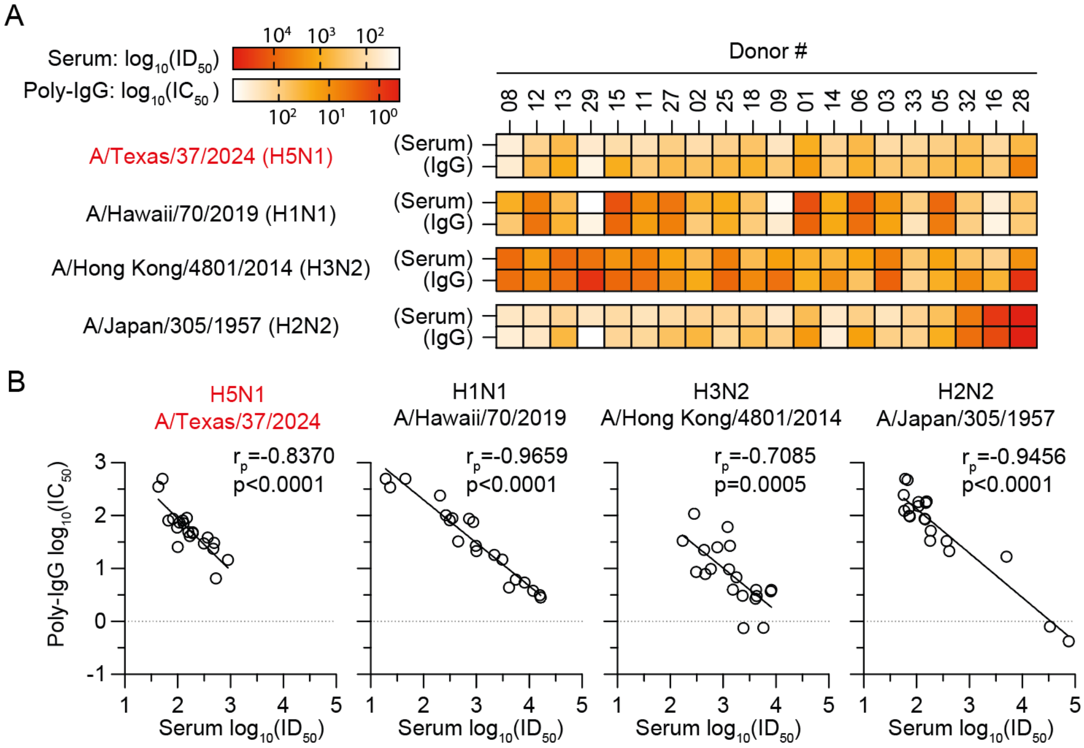


**Figure S4. Comparison of serum vs. plasma-derived poly-IgG neutralization, related to Figure 1**. (**A**) Sera and purified plasma IgG dilutions from the same 20 individuals were tested for HA-pseudotype lentivirus neutralization against the indicated strains and half-maximal inhibitory dilutions ($ID_{50}$) or concentrations ($IC_{50}$) values were determined. (**B**) Correlation of $log_{10}(IC_{50})$ and $log_{10}(ID_{50})$ from (A). Pearson correlation coefficients ($r_p$) and corresponding p-values are given in the plots. The line represents a linear regression of the data

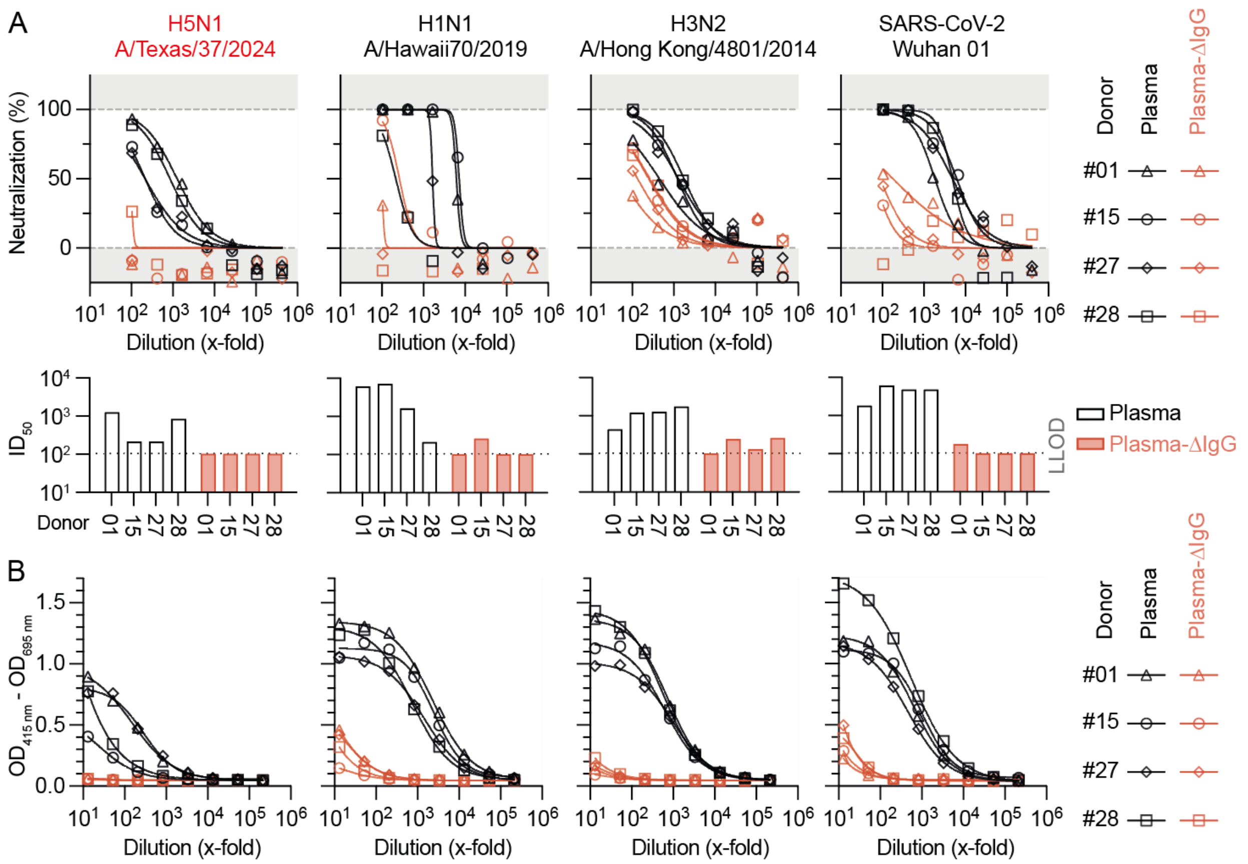


**Figure S5. HA-pseudotyped virus neutralization and HA IgG binding before and after IgG depletion from plasma, related to Figure 1.** (**A**) Plasma samples from n = 4 donors were incubated with Protein A/G sepharose to deplete poly-IgGs and were assessed for neutralization activity against pseudotyped lentivirus expressing the indicated hemagglutinins (H5, H1, H3) or SARS-CoV-2 spike protein (Wuhan 01) as a control. Symbols depict normalized neutralization from n = 2 technical replicates for the different donors with lines representing non-linear curve fits. Bar graphs show calculated half-maximal inhibitory dilutions ($ID_{50}$, x-fold dilution) from non-linear curve fits for each individual (1-4) before (white bars) and after (red bars) depletion. Lower limit of detection (LLOD) was 100-fold dilution. (**B**) Poly-IgG depleted and non-depleted plasma from (A) was tested for IgG reactivity against recombinant HAs from the indicated hemagglutinins (H5, H1, H3) or SARS-CoV-2 spike protein (Wuhan 01) as a positive control by ELISA. Curves represent non-linear fits of $OD_{415\ nm}$ - $OD_{695\ nm}$ from n = 2 technical replicates. LLOD for ELISAs was 13-fold.

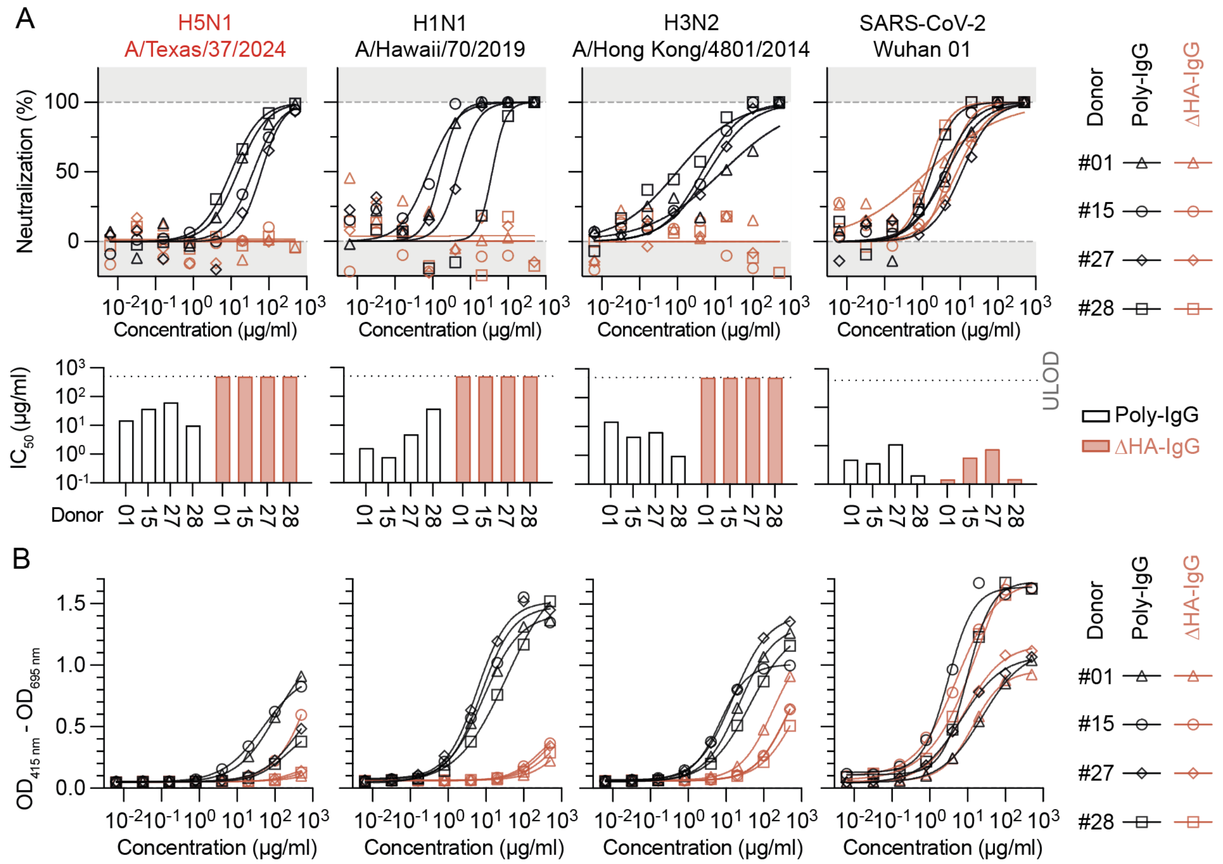


**Figure S6. HA-pseudotyped virus neutralization and HA binding before and after depletion of HA-reactive antibodies from poly-IgG, related to Figure 1.** (**A**) Purified poly-IgG samples from n = 4 donors were incubated with immobilized HAs from H5N1 (A/Texas/37/2024), H1N1 (A/Hawaii/70/2019) and H3N2 (A/Hong Kong/4801/2014) to deplete HA-reactive antibodies (ΔHA-IgG) and assessed for neutralization activity against pseudotyped lentivirus expressing the indicated hemagglutinins (H5, H1, H3) or SARS-CoV-2 spike protein (Wuhan 01) as a negative control. Symbols depict normalized neutralization from n = 2 technical replicates for the different donors with lines representing non-linear curve fits. Bar graphs show calculated half-maximal inhibitory concentration (IC50) values from non-linear curve fits for each individual (1-4) before (white bars) and after (red bars) depletion. Upper limit of detection (ULOD) was 500 µg/ml. (**B**) HA-Ab depleted and non-depleted poly-IgGs from (A) were tested for reactivity against recombinant HAs from the indicated hemagglutinins (H5, H1, H3) or SARS-CoV-2 spike protein (Wuhan 01) as a negative control by ELISA. Curves represent non-linear fits of $OD_{415\ nm}$ - $OD_{695\ nm}$ from n = 2 technical replicates.

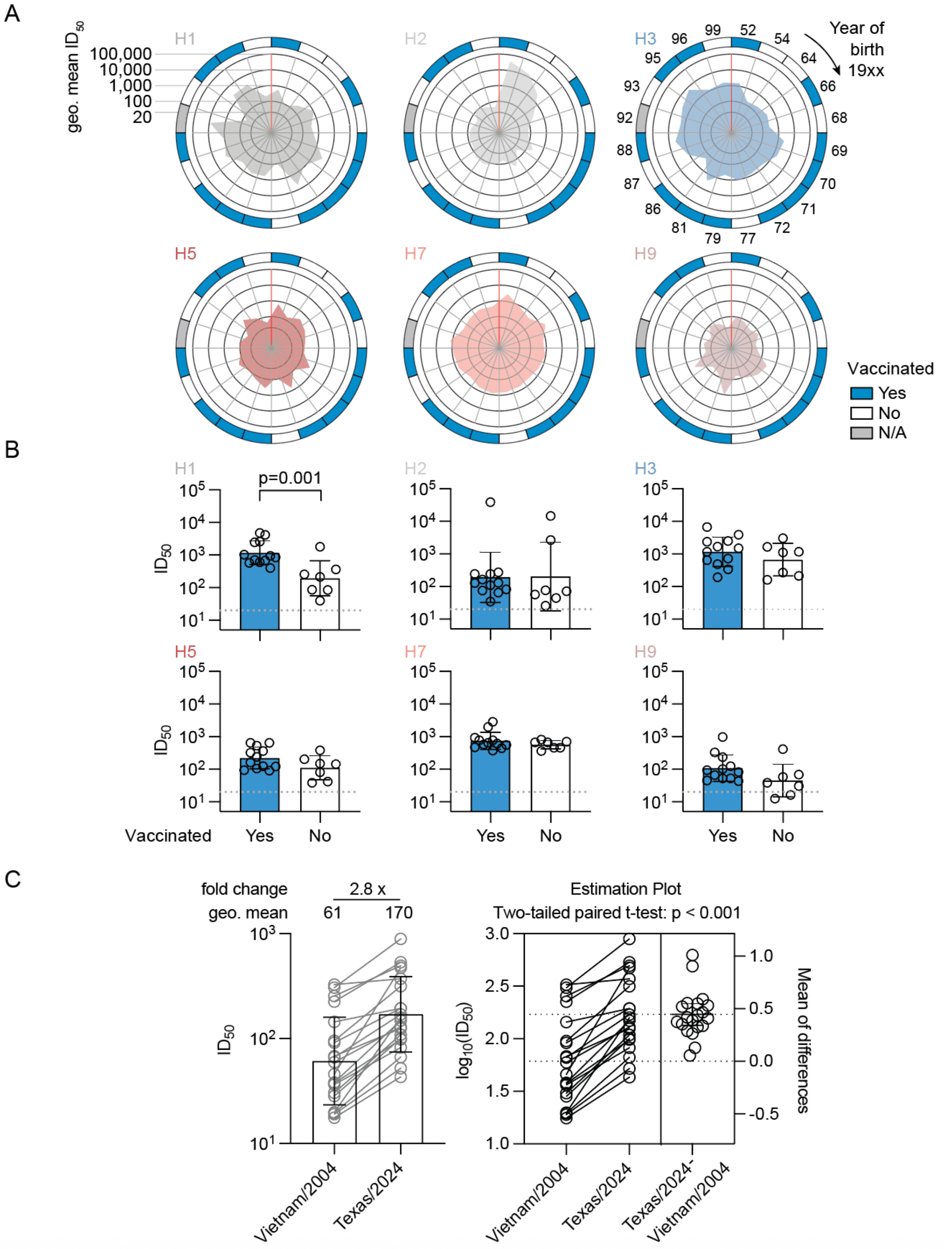


**Figure S7. HA subtype neutralization per individual, related to Figure 3.** (**A**) Geometric mean half-maximal inhibitory dilution ($ID_{50}$) was calculated for each of n = 20

individuals over all strains included in the depicted subtypes (i.e., n = 15 H1, n = 2 H2, n = 40 H3, n = 12 H5, n = 4 H7, and n = 4 H9). Each slice of the pie charts represents one individual and individuals were sorted clockwise by their year of birth starting from the oldest participant born in 1952. Calculated values for each individual are plotted within the center of each pie. (**B**) Geometrc mean $ID_{50}$ values as in (A) stratified by self-reported vaccination status. Bar graphs depict geometric mean values with geometric mean standard deviations as error bars. The individual that did not report any vaccination information was excluded from the analysis. $ID_{50}$ values were arbitrarily set to 10, if they could not be inferred by curve fits or where >2 fold lower than the lower limit of detection (i.e., inferred $ID_{50}<10$). P values are depicted if ≤0.05 and have been determined by a two-tailed t test. (**C**) Individual $ID_{50}$ values (as in main Figure 3B) for A/Vietnam/1203/2004 and A/Texas/37/2024 are plotted as linked dots. Bars in the background represent the geometric mean $ID_{50}$ values with geometric standard deviations as error bars (left panel). A two-tailed paired t-test was performed on the $log_{10}$-transformed $ID_{50}$ values to test for significance (right panel).

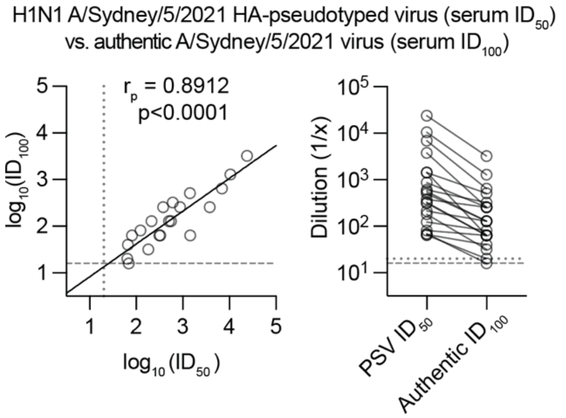


**Figure S8. Correlation of $ID_{50}$ and $ID_{100}$ values from H1N1 pseudovirus and authentic virus neutralization assay, related to Figure 3.** Comparison of serum neutralization (n=20) for H1N1 A/Sydney/5/2021 HA-pseudotyped lentivirus ($ID_{50}$, data from Figure 3B) and authentic H1N1 A/Sydney/5/2021 ($ID_{100}$). The pearson correlation coefficient $r_p$ and the corresponding p-value for the $log_{10}$-transformed data is given in the figure. Dashed/dotted lines represent the LLOD for the authentic virus and pseudovirus neutralization tests, respectively.

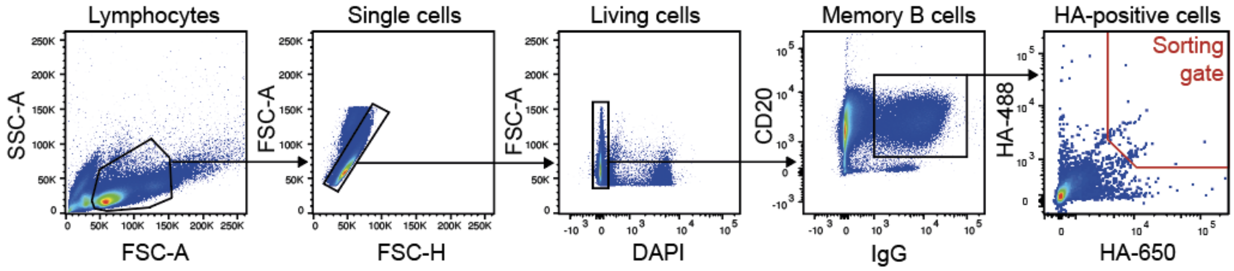


**Figure S9. Example gating strategy for sorting of H5N1 A/Texas/37/2024 HA(Y98F)-specific memory B cells, related to Figure 4.** H5N1 A/Texas/37/2024 HA(Y98F)-specific memory B cells were purified from CD19-enriched PBMCs through flow-cytometry by gating on lymphocytes in forward and side scatter (FSC-A, SSC-A), exclusion of doublets through forward scatter height (FSC-H) versus area (FSC-A) discrimination, exclusion of DAPI positive (i.e., dead) cells, selection of CD20 positive and IgG positive cells, and selection of H5N1 A/Texas/37/2024 HA(Y98F)-DyLight488 and -Dylight650 double positive cells.

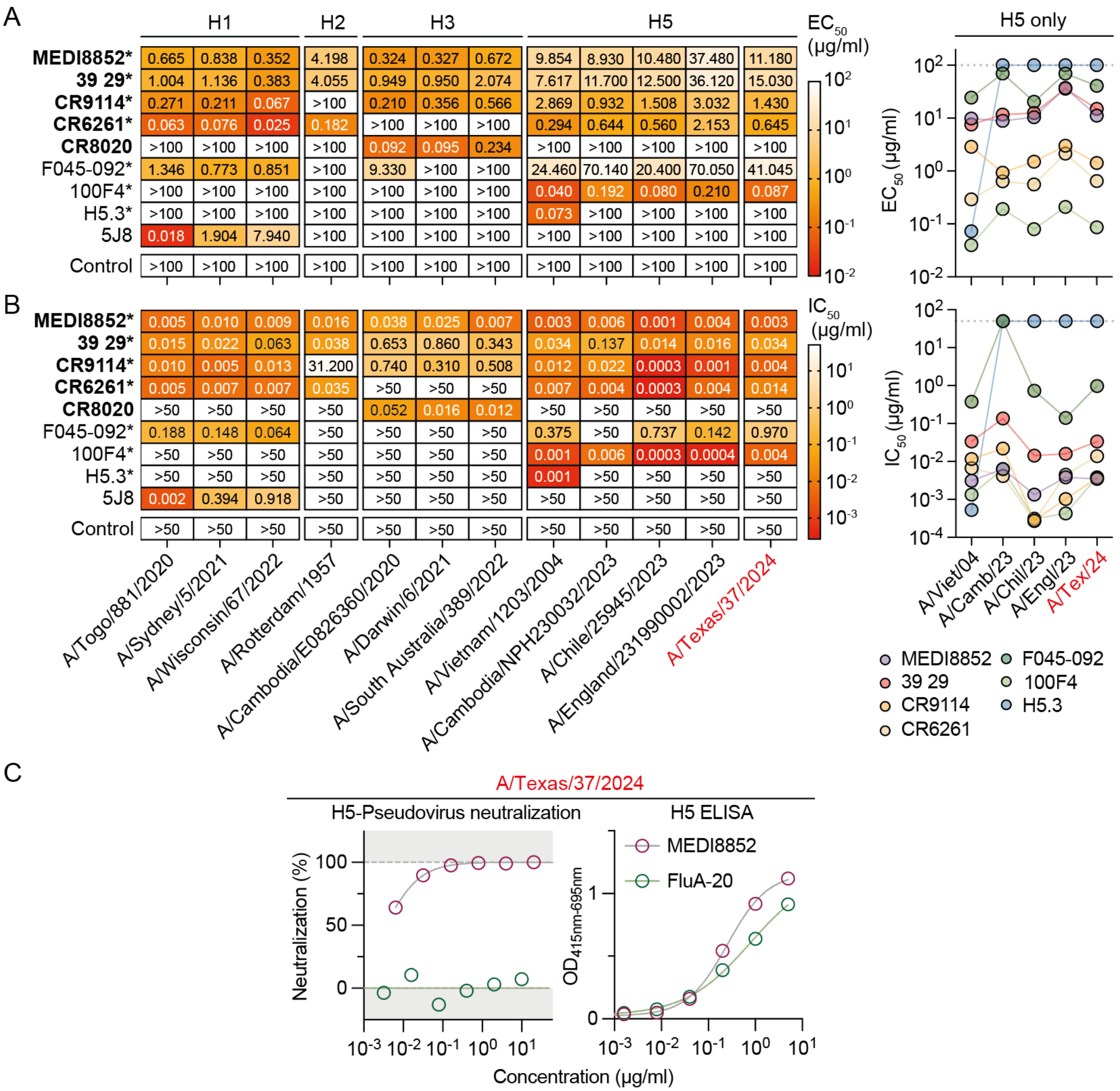


**Figure S10. H5-reactive and clinically developed monoclonal antibodies bind and neutralize A/Texas/37/2024 HA, related to Figure 5.** (**A**) Half-maximal effective concentration ($EC_{50}$) of n = 7 H5-reactive as well as two H1- or H3-specific monoclonal antibodies determined by cell-surface-expressed HA binding against 12 different HA variants. The dot-plot on the right shows $EC_{50}$-values from the n = 7 H5-reactive mAbs against H5 virus strains only. (**B**) Heatmap and dot-plot of half-maximal inhibitory concentrations ($IC_{50}$) of the same monoclonal antibodies as in (A) determined by neutralization tests against HA-pseudotyped lentiviruses expressing the same HA variants as in (A). Antibodies in clinical trials are highlighted in bold. Control: SARS-CoV-2 antibody CnC2t1p1_D6. *Pre-described H5-reactive mAbs. (**C**) HA A/Texas/37/2024 pseudovirus neutralization and binding (ELISA) data for the H5-reactive non-neutralizing antibody FluA-20 in comparison to MEDI8852.

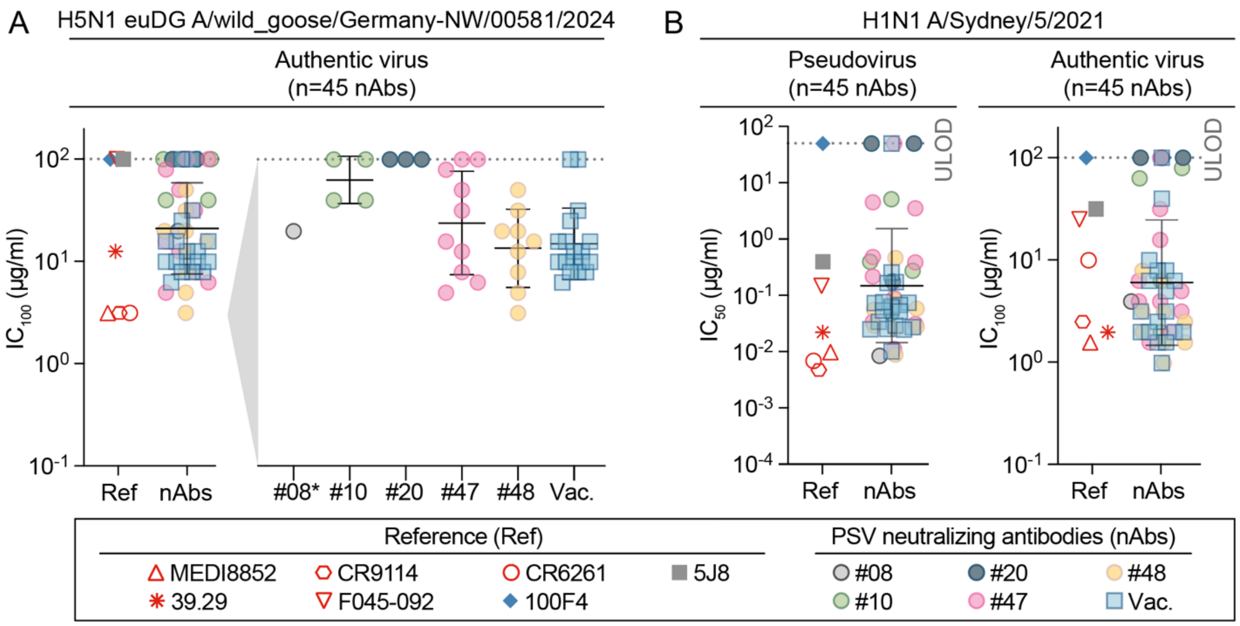


**Figure S11. H5N1 (euDG) and H1N1 authentic virus as well as H1N1 HA-pseudotyped virus neutralization by monoclonal antibodies, related to Figure 5.** (**A**) Complete (100%) inhibitory concentrations ($IC_{100}$) of n = 45 H5-neutralizing antibodies (nAbs) and n = 7 reference antibodies as in Figure 5, determined by microneutralization test against authentic highly pathogenic avian influenza virus (A/wild goose/Germany-NW/2024AI02730/2024, genotype euDG) (**B**) Half-maximal inhibitory concentration ($IC_{50}$) and $IC_{100}$ of the same antibodies as in (A), determined by neutralization assay against pseudotyped lentivirus expressing the HA of A/Sydney/5/2021 (left plot) or determined by microneutralization test against authentic A/Sydney/5/2021 (H1N1 Sydney/21). Mean values and error bars in depict geometric mean and geometric standard deviation. Dashed lines represent the upper limit of detection (ULOD) with 50 µg/ml and 100 µg/ml for pseudotyped and authentic virus, respectively.

**Table S1.** Blood donor information, related to Figures 1-5, S3-7, and S11

| ID | Sex | Year of birth | Sampling date | Age at sampling | Any Influenza vaccination | Last vaccination date | Known prior Influenza A infection | Last known infection date |
|---|---|---|---|---|---|---|---|---|
| 01 | F | 1972 | Aug-23 | 51 | Yes | Oct-21 | No | N/A |
| 02 | F | 1986 | Aug-23 | 36 | Yes | Oct-21 | No | N/A |
| 03 | F | 1969 | Aug-23 | 54 | Yes | Oct-22 | No | N/A |
| 04 | F | 1989 | Aug-23 | 33 | No | N/A | No | N/A |
| 05 | F | 1966 | Aug-23 | 56 | Yes | Nov-20 | No | N/A |
| 06 | M | 1970 | Aug-23 | 53 | Yes | Dec-22 | No | N/A |
| 07 | F | 1972 | Aug-23 | 50 | Yes | Oct-09 | No | N/A |
| 08 | F | 1999 | Aug-23 | 24 | No | N/A | Yes | 2015 |
| 09 | M | 1977 | Aug-23 | 46 | No | N/A | No | N/A |
| 10 | M | 1967 | Aug-23 | 55 | No | N/A | No | N/A |
| 11 | F | 1988 | Aug-23 | 34 | Yes | Nov-22 | No | N/A |
| 12 | F | 1996 | Aug-23 | 26 | Yes | Oct-21 | No | N/A |
| 13 | F | 1995 | Aug-23 | 28 | Yes | Oct-22 | No | N/A |
| 14 | M | 1971 | Aug-23 | 51 | Yes | Oct-22 | No | N/A |
| 15 | M | 1992 | Aug-23 | 31 | N/A | N/A | No | N/A |
| 16 | F | 1954 | Aug-23 | 69 | No | N/A | No | N/A |
| 17 | F | 1961 | Aug-23 | 61 | Yes | Jul-05 | No | N/A |
| 18 | F | 1979 | Aug-23 | 44 | Yes | Sep-22 | No | N/A |
| 19 | M | 1969 | Aug-23 | 53 | Yes | Nov-22 | No | N/A |
| 20 | M | 1966 | Aug-23 | 57 | Yes | Nov-22 | No | N/A |
| 21 | M | 1987 | Aug-23 | 36 | Yes | Dec-22 | No | N/A |
| 22 | F | 1995 | Aug-23 | 27 | Yes | Feb-20 | No | N/A |
| 23 | F | 1969 | Aug-23 | 54 | Yes | Nov-12 | No | N/A |
| 24 | F | 1989 | Aug-23 | 34 | Yes | Oct-22 | No | N/A |
| 25 | M | 1981 | Aug-23 | 42 | Yes | Oct-22 | No | N/A |
| 26 | F | 1977 | Aug-23 | 46 | No | N/A | No | N/A |
| 27 | M | 1987 | Aug-23 | 35 | No | N/A | No | N/A |
| 28 | F | 1952 | Aug-23 | 71 | Yes | N/A | Yes | N/A |
| 29 | M | 1993 | Aug-23 | 30 | No | N/A | No | N/A |
| 30 | F | 1997 | Sep-23 | 26 | Yes | Jul-05 | No | N/A |
| 31 | F | 1972 | Sep-23 | 51 | Yes | Oct-22 | No | N/A |
| 32 | M | 1964 | Sep-23 | 59 | No | N/A | No | N/A |
| 33 | M | 1968 | Sep-23 | 54 | No | N/A | No | N/A |
| 34 | F | 1996 | Nov-23 | 27 | No | N/A | N/A | N/A |
| 35 | F | 1984 | Nov-23 | 39 | Yes | Oct-23 | No | N/A |
| 36 | M | 1952 | Nov-23 | 71 | No | N/A | No | N/A |
| 37 | F | 2004 | Dec-23 | 19 | No | N/A | No | N/A |
| 38 | F | 1978 | Dec-23 | 45 | Yes | Oct-23 | No | N/A |
| 39 | F | 1994 | Jan-24 | 29 | No | N/A | No | N/A |
| 40 | F | 1966 | Apr-24 | 57 | Yes | Sep-06 | No | N/A |
| 41 | F | 1971 | Apr-24 | 52 | Yes | >15 years | No | N/A |
| 42 | M | 1966 | Apr-24 | 58 | Yes | >10 years | No | N/A |
| 43 | M | 1965 | Apr-24 | 58 | Yes | Sep-23 | No | N/A |
| 44 | M | 1997 | Apr-24 | 26 | Yes | Nov-21 | No | N/A |
| 45 | F | 1966 | Apr-24 | 57 | Yes | Oct-23 | No | N/A |
| 46 | F | 1996 | Apr-24 | 27 | Yes | Oct-21 | No | N/A |
| 47 | F | 1986 | May-24 | 38 | No | N/A | Yes | Feb-24 |
| 48 | F | 1996 | Jun-24 | 28 | Yes | 2023 | No | N/A |
| 49 | M | 1998 | Jun-24 | 26 | No | N/A | No | N/A |
| 50 | F | 1989 | Jun-24 | 34 | Yes | Jan-24 | Yes | 2019 |
| 51 | F | 2000 | Jun-24 | 23 | No | N/A | No | N/A |
| 52 | F | 1999 | Jun-24 | 24 | Yes | Nov-23 | No | N/A |
| 53 | F | 1994 | Jun-24 | 29 | Yes | N/A | No | N/A |
| 54 | F | 1986 | Jun-24 | 37 | Yes | Nov-23 | No | N/A |
| 55 | F | 1987 | Jun-24 | 36 | Yes | Nov-23 | No | N/A |
| 56 | F | 2001 | Jun-24 | 22 | Yes | Nov-23 | No | N/A |
| 57 | M | 1986 | Jun-24 | 38 | Yes | Oct-23 | No | N/A |
| 58 | F | 1998 | Jun-24 | 25 | Yes | Oct-23 | No | N/A |
| 59 | M | 1993 | Jun-24 | 30 | Yes | Feb-23 | No | N/A |
| 60 | F | 1981 | Jun-24 | 43 | Yes | Oct-22 | No | N/A |
| 61 | F | 1963 | Jun-24 | 60 | No | N/A | No | N/A |
| 62 | M | 1975 | Jun-24 | 49 | Yes | Nov-23 | Yes | N/A |
| 63 | F | 1980 | Jun-24 | 43 | Yes | Nov-22 | No | N/A |
| 64 | F | 1995 | Jun-24 | 29 | Yes | Nov-23 | Yes | Jan-23 |
| 65 | F | 1994 | Jun-24 | 29 | No | N/A | No | N/A |
| 66 | F | 2002 | Jun-24 | 22 | Yes | Oct-22 | Yes | 2010 |

**Table S2.** Influenza strain information, related to Figures 1, 3, S3, and S10

| # | HxNy | Strain | EPI |
|---|---|---|---|
| 1 | H1N1 | A/Wisconsin/67/2022 | EPI_ISL_15928538 |
| 2 | H1N1 | A/Sydney/5/2021 | EPI_ISL_6424984 |
| 3 | H1N1 | A/Togo/881/2020 | EPI_ISL_1015524 |
| 4 | H1N1 | A/Ghana/140/2020 | EPI_ISL_959864 |
| 5 | H1N1 | A/Perth/41/2020 | EPI_ISL_482750 |
| 6 | H1N1 | A/Victoria/2570/2019 | EPI_ISL_548964 |
| 7 | H1N1 | A/Wisconsin/588/2019 | EPI_ISL_404460 |
| 8 | H1N1 | A/Guangdong-Maonan/SWL1536/2019_C1C1 | EPI_ISL_391019 |
| 9 | H1N1 | A/Guangdong-Maonan/SWL1536/2019_E4 | EPI_ISL_392654 |
| 10 | H1N1 | A/Hawaii/70/2019 | EPI_ISL_400916 |
| 11 | H1N1 | A/Brisbane/64/2019 | EPI_ISL_389532 |
| 12 | H1N1 | A/Oman/5442/2018 | EPI_ISL_347417 |
| 13 | H1N1 | A/Brisbane/2/2018 | EPI_ISL_410633 |
| 14 | H1N1 | A/Brisbane/59/2007 | EPI_ISL_23380 |
| 15 | H1N1 | A/Beijing/262/1995 | EPI_ISL_22625 |
| 16 | H2N2 | A/Rotterdam/1957 | EPI_ISL_84911 |
| 17 | H2N2 | A/Japan/305/1957 | EPI_ISL_5869 |
| 18 | H3N2 | A/South Australia/389/2022 | EPI_ISL_16613678 |
| 19 | H3N2 | A/Darwin/6/2021 | EPI_ISL_3534319 |
| 20 | H3N2 | A/Darwin/9/2021 | EPI_ISL_3100998 |
| 21 | H3N2 | A/Cambodia/E0826360/2020 | EPI_ISL_1296487 |
| 22 | H3N2 | A/Bangladesh/1813/2020 | EPI_ISL_882827 |
| 23 | H3N2 | A/Sydney/473/2019 | EPI_ISL_406676 |
| 24 | H3N2 | A/Ghana/3532/2019 | EPI_ISL_406176 |
| 25 | H3N2 | A/Kanagawa/159/2019 | EPI_ISL_398727 |
| 26 | H3N2 | A/Hong Kong/2671/2019_E7 | EPI_ISL_400888 |
| 27 | H3N2 | A/Hong Kong/2671/2019_C1S2 | EPI_ISL_391201 |
| 28 | H3N2 | A/South Australia/34/2019_E4 | EPI_ISL_400538 |
| 29 | H3N2 | A/South Australia/34/2019_SIAT1 | EPI_ISL_346283 |
| 30 | H3N2 | A/Sydney/549/2017 | EPI_ISL_396727 |
| 31 | H3N2 | A/Victoria/659/2017 | EPI_ISL_396885 |
| 32 | H3N2 | A/Victoria/601/2017 | EPI_ISL_396870 |
| 33 | H3N2 | A/Victoria/723/2017 | EPI_ISL_289904 |
| 34 | H3N2 | A/Tunisia/1597/2017 | EPI_ISL_262280 |
| 35 | H3N2 | A/Victoria/2030/2017 | EPI_ISL_396820 |
| 36 | H3N2 | A/Victoria/2014/2017 | EPI_ISL_271306 |
| 37 | H3N2 | A/Kansas/14/2017_E17 | EPI_ISL_350966 |
| 38 | H3N2 | A/Kansas/14/2017_S2 | EPI_ISL_292575 |
| 39 | H3N2 | A/Switzerland/8060/2017 | EPI_ISL_294252 |
| 40 | H3N2 | A/Canberra/7/2016 | EPI_ISL_227639 |
| 41 | H3N2 | A/Singapore/INFIMH160019/2016 | EPI_ISL_285898 |
| 42 | H3N2 | A/South Australia/118/2016 | EPI_ISL_239826 |
| 43 | H3N2 | A/Hong Kong/4801/2014 | EPI_ISL_165554 |
| 44 | H3N2 | A/Switzerland/9715293/2013 | EPI_ISL_162149 |
| 45 | H3N2 | A/Texas/50/2012 | EPI_ISL_122006 |
| 46 | H3N2 | A/Victoria/361/2011 | EPI_ISL_134450 |
| 47 | H3N2 | A/Perth/16/2009 | EPI_ISL_31055 |
| 48 | H3N2 | A/Brisbane/10/2007 | EPI_ISL_23194 |
| 49 | H3N2 | A/Wisconsin/67/2005 | EPI_ISL_115646 |
| 50 | H3N2 | A/California/7/2004 | EPI_ISL_113070 |
| 51 | H3N2 | A/Moscow/10/1999 | EPI_ISL_127595 |
| 52 | H3N2 | A/Sydney/5/1997 | EPI_ISL_29667 |
| 53 | H3N2 | A/Nanchang/933/1995 | EPI_ISL_111174 |
| 54 | H3N2 | A/Netherlands/179/1993 | EPI_ISL_111045 |
| 55 | H3N2 | A/Netherlands/620/1989 | EPI_ISL_114347 |
| 56 | H3N2 | A/Bilthoven/1761/1976 | EPI_ISL_113970 |
| 57 | H3N2 | A/Bilthoven/23337/1972 | EPI_ISL_110819 |
| 58 | H5N1 | A/Texas/37/2024 | EPI_ISL_19027114 |
| 59 | H5N1 | A/England/231990002/2023 | EPI_ISL_17736649 |
| 60 | H5N1 | A/Chile/25945/2023 | EPI_ISL_17468386 |
| 61 | H5N1 | A/Cambodia/NPH230032/2023 | EPI_ISL_17024123 |
| 62 | H5N8 | A/Astrakhan/3212/2020 | EPI_ISL_1038924 |
| 63 | H5N6 | A/Hubei/29578/2016 | EPI_ISL_256213 |
| 64 | H5N6 | A/Guangxi/55726/2016 | EPI_ISL_240704 |
| 65 | H5N1 | A/Egypt/N0005/2015 | EPI_ISL_17767282 |
| 66 | H5N1 | A/Indonesia/NIHRD15028/2015 | EPI_ISL_195732 |
| 67 | H5N1 | A/Cambodia/X0123311/2013 | EPI_ISL_166636 |
| 68 | H5N1 | A/Vietnam/1203/2004 | EPI_ISL_10656749 |
| 69 | H5N1 | A/Hong Kong/213/03 | EPI_ISL_268 |
| 70 | H7N9 | A/Hebei/27401/2017 | EPI_ISL_285159 |
| 71 | H7N9 | A/Anhui/01887/2014 | EPI_ISL_285159 |
| 72 | H7N7 | A/Italy/3/2013 | EPI_ISL_171318* |
| 73 | H7N1 | A/Shangdong/2/2013 | EPI_ISL_17264959 |
| 74 | H9N2 | A/Oman/2747/2019 | EPI_ISL_353983 |
| 75 | H9N2 | A/Anhui-Lujiang/39/2018 | EPI_ISL_330737 |
| 76 | H9N2 | A/Bangladesh/0994/2011 | EPI_ISL_140388 |
| 77 | H9N2 | A/Chicken/Hong Kong/G9/1997 | EPI_ISL_146698 |

* Sequence was corrected after alignment

**Table S3.** Monoclonal antibodies, related to Figures 5, S10 and S11

| # | Antibody name | Origin | Target | In clinical development | Reference |
|---|---|---|---|---|---|
| 1 | MEDI8852 | Blood donor in switzerland; genetically optmized version of FY1 | HA stalk | Yes, VIR2482, NCT02350751, NCT02603952 | Kallewaard et al. Cell, 2016 |
| 2 | 39.29 | 2009 seasonal influenza vaccine recipient | HA stalk | Yes, Gedivumab (MHAA4549A); NCT01877785, NCT02284607, NCT01980966, NCT02293863, NCT02623322 | Nakamura et al. Cell Host Microbe, 2013 |
| 3 | CR9114 | Seasonal influenza vaccine recipient | HA stalk | Yes, CTIS2023-505020-57-00 | Dreyfus et al. Science, 2012 |
| 4 | CR6261 | Seasonal influenza vaccine recipient | HA stalk | Yes; NCT02371668, NCT01406418 | Throsby et al. PLoS One, 2008 |
| 5 | F045-092 | Apharesis donor, born in 1974 | HA head | No | Ohshima et al. J Virol, 2011 |
| 6 | 100F4 | H5N1 subclade 2.3.4 convalenscent donor | HA head | No | Hu et al. J Virol, 2012 |
| 7 | H5.3 | Participants of a H5N1 (A/Vietnam/1203/2004) vaccine trial | HA head | No | Thornburg et al. J Clin Invest, 2013 |
| 8 | 5J8 | Middle-aged woman | HA head | No | Krause et al. J Virol, 2011 |
| 9 | CR8020 | 2006/2007 Seasonal influenza vaccinated donors | HA stalk | Yes; NCT01938352, NCT01756950 | Ekiert et al. Science, 2011 |
| 10 | CnC2t1p1_D6 | SARS-CoV-2 convalescent donor | SARS-CoV-2 RBD | No | Kreer et al. Cell, 2020 |
| 11 | DZIF-10c (HbnC3t1p1_F4) | SARS-CoV-2 convalescent donor | SARS-CoV-2 RBD | Yes; EudraCT Number:2020-004448-27 | Kreer et al. Cell, 2020; Halwe et al. Viruses, 2021 |
| 12 | FluA-20 | Clinical trial particpant of experimental H5N1 and H7N9 vaccines | HA head | No | Bangaru et al., 2019 |
